# Multiple genes *in cis* mediate the effects of a single chromatin accessibility variant on aberrant synaptic development and function in human neurons

**DOI:** 10.1101/2021.12.11.472229

**Authors:** Siwei Zhang, Hanwen Zhang, Marc P. Forrest, Yifan Zhou, Vikram A. Bagchi, Alena Kozlova, Marc Dos Santos, Nicolas H. Piguel, Leonardo E. Dionisio, Alan R. Sanders, Zhiping P. Pang, Xin He, Peter Penzes, Jubao Duan

## Abstract

Despite hundreds of risk loci from genome-wide association studies of neuropsychiatric disorders, causal variants/genes remain largely unknown. Here, in *NEUROG2*-induced human neurons, we identified 31 risk SNPs in 26 schizophrenia (SZ) risk loci that displayed allele-specific open chromatin (ASoC) and were likely to be functional. Editing the strongest ASoC SNP rs2027349 near *vacuolar protein sorting 45 homolog* (*VPS45*) altered the expression of *VPS45*, lncRNA *AC244033.2*, and a distal gene, *C1orf54*, in human neurons. Notably, the global gene expression changes in neurons were enriched for SZ risk and correlated with post-mortem brain gene expression signatures of neuropsychiatric disorders. Neurons carrying the risk allele exhibited increased dendritic complexity, synaptic puncta density, and hyperactivity, which were reversed by knocking-down distinct *cis*-regulated genes (*VPS45*, *AC244033.2*, or *C1orf54*), suggesting a phenotypic contribution from all three genes. Interestingly, transcriptomic analysis of knockdown cells suggested a non-additive effects of these genes. Our study reveals a compound effect of multiple genes at a single SZ locus on synaptic development and function, providing a mechanistic link between a non-coding SZ risk variant and disease-related cellular phenotypes.

## Introduction

Genome-wide association studies (GWAS) of schizophrenia (SZ) and other neuropsychiatric disorders have identified hundreds of common genetic risk loci ^1–5^. Most neuropsychiatric risk variants reside in noncoding sequences that likely alter gene expression. However, despite the progress made from studying human postmortem brains (*e.g.*, PsychENCODE) ^6–9^, animal models ^10^, and, more recently, human induced pluripotent stem cell (hiPSC) models ^11–14^, causal disease genes and the underlying molecular mechanisms remain elusive for most neuropsychiatric disease risk loci.

A challenge for understanding the causal mechanisms underlying these associations is that each GWAS locus often encompasses multiple genes and many common risk variants that are equally associated with disease due to linkage disequilibrium (LD). Co-localization analysis of disease risk variants with regulatory DNA regions defined by postmortem brain epigenomic marks, *e.g.*, PsychENCODE ^15^, GTEx ^16^, and brain expression QTL (eQTL) ^17^, has been commonly used to prioritize putatively functional variants. These studies, however, have significant limitations: (1) regulatory regions are often only modestly enriched for disease variants, presumably because most variants in these regions are not functional; (2) QTL do not directly point to causal variants because of LD; and (3) most of the existing work was performed on adult brain samples, which may miss regulatory variants of neuropsychiatric disorders that are often early developmental stage- and cell type-specific ^18^. As a result, for most neuropsychiatric risk loci, the specific causal variants, and their regulated risk genes, as well as disease-relevant cellular phenotypes, remain largely unidentified.

We have recently shown that allele-specific open chromatin (ASoC) is an effective functional readout of regulatory single nucleotide polymorphisms (SNPs), displaying differential allelic chromatin accessibility in heterozygous individuals ^13^, *i.e.*, two alleles of an SNP show read imbalance in Assay for Transposase-Accessible Chromatin using sequencing (ATAC-seq) ^13, 19^. Compared to eQTL mapping, ASoC mapping has the advantage of directly identifying putatively functional variants rather than those merely in LD ^13^. ASoC SNPs are often cell type-specific, and neuronal ASoC SNPs are strongly enriched for neuropsychiatric risk variants ^13^. Here, we systematically identified the putatively functional SZ GWAS risk variants ^1^ that showed ASoC in hiPSC-derived and *NEUROG2* (NGN2)-induced excitatory neurons (NGN2-Glut) ^20^, a widely used neuronal model for characterizing neuropsychiatric cellular phenotypes ^11, 21–24^. Combined with results from a multiplex CRISPRi screening for *cis*-target genes of ASoC sequences, we prioritized the SZ-associated ASoC SNP, rs2027349, near *vacuolar protein sorting 45 homolog* (*VPS45)* for functional characterization in NGN2-Glut. We found that this single SZ risk SNP *cis*-regulates two adjacent genes, *VPS45* and a long noncoding RNA (lncRNA, *AC244033.2*), as well as a distal gene that encodes *chromosome 1 open reading frame 54* (*C1orf54*). All three genes synergistically contributed to the cellular and molecular alterations relevant to neuropsychiatric disorders.

## Results

### ASoC mapping in NGN2-Glut identifies functional SZ risk SNPs

To identify functional SZ GWAS risk variants that may alter chromatin accessibility and gene expression in NGN2-Glut, we carried out ATAC-seq ^19^ and mapped ASoC variants in neurons of 20 hiPSC lines (Fig. 1a, Extended Data Fig. 1a-b; Methods). We obtained 48∼66 M paired-end ATAC-seq reads per sample (Supplementary Table 1, Extended Data Fig. 1c-f). With 183,692 open chromatin peaks (accessible under GSE188491) of NGN2-Glut called by using MACS2 ^25^ at a false discovery rate (FDR) < 0.05, we compared their similarity to our previously generated open chromatin profiles of neurons at a different cellular state, *i.e.*, day-15 glutamatergic neurons directly induced from hiPSC-derived neural progenitor cells (iN-Glut) ^13^ (Fig. 1a). We found that although NGN2-Glut and iN-Glut samples were overall similar to each other, they did display different open chromatin profiles between NGN2-Glut and iN-Glut (Extended Data Fig. 1g).

**Fig. 1:**
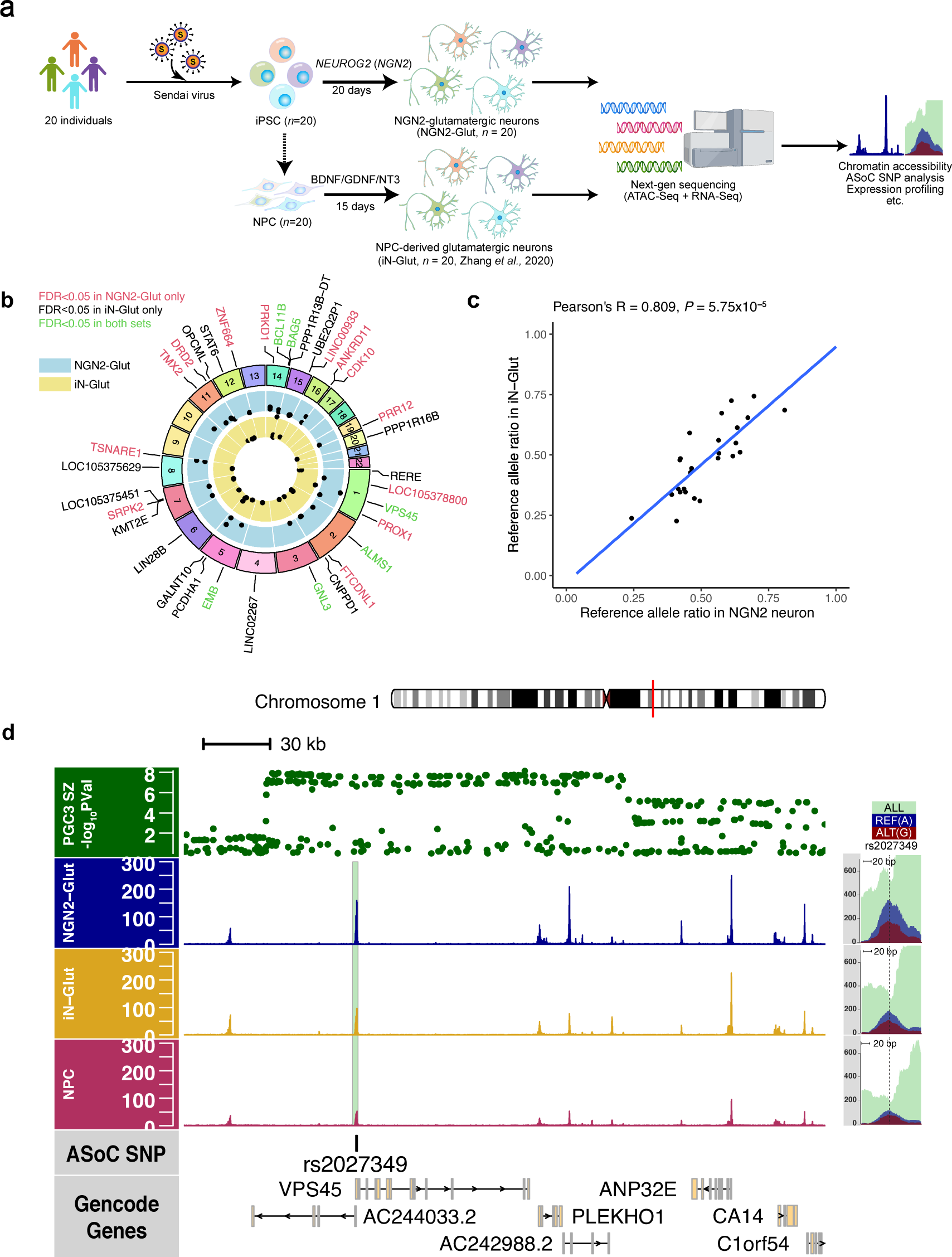
Allele-specific open chromatin (ASoC) mapping in NGN2-Glut identifies strong ASoC at the SZ-associated *VPS45* locus. (a) Schematics depicting the strategy to generate NGN2-induced, in comparison with previously NPC-derived (dashed arrows), glutamatergic excitatory neuron from hiPSC for bulk ATAC-seq and RNA-seq analysis. (b) Circus plot showing SZ-credible risk SNPs that exhibited ASoC (FDR < 0.05) in NGN2-Glut, iN-Glut, or both. (c) Allelic ratio correlation of SZ-associated ASoC SNPs (FDR < 0.05) in iN-Glut-15 and NGN2-Glut, *n* = 26. (d) Cell type-specific and allele-specific ATAC-seq read pileups flanking ASoC SNP rs2027349 at the SZ-associated *VPS45* locus. The top track shows PGC3 SZ GWAS *p*-values of all the SNPs in this region. Aquamarine: total reads pileup for rs2027349; dark blue: reads pileup containing reference allele (G); dark red: reads pileup containing alternative allele (A).

We identified 8,205 ASoC SNPs (FDR < 0.05) out of 163,066 heterozygous SNPs in open chromatin regions (OCRs) (Extended Data Fig. 2a and Supplementary Table 2). Since we have previously characterized the cell type-specificity, transcriptional factor binding site (TFBS) disruption, and regulatory properties of ASoC SNPs ^13^, here we focused on identifying putatively functional SZ GWAS risk SNPs ^26^ showing ASoC in NGN2-Glut, in comparison with those previously identified in iN-Glut (Supplementary Table 3) ^13^. For the list of SZ credible SNPs at the 108 risk loci (PGC2 SZ GWAS) ^27^, we found 31 ASoC SNPs at 26 SZ risk loci in NGN2-Glut, of which 14 were heterozygous and 7 of them also showed ASoC in iN-Glut (Fig. 1b and Supplementary Table 3). Of the 17 SZ loci with genome-wide significant (GWS) risk SNPs showing ASoC in iN-Glut ^13^, 9 had a nominally significant difference of ASoC (*p*-value < 0.05) in NGN2-Glut. Overall, there was a significant overlap of SZ risk SNPs showing ASoC (FDR < 0.05) between the two datasets (24-fold enrichment, *p*-value < 2.5 × 10^-6^; 2-tailed Fisher’s Exact Test) (Extended Data Fig. 2b, Supplementary Table 3). The allelic ratio of the chromatin accessibility of all SZ credible SNPs also showed a modest correlation between NGN2-Glut and iN-Glut (*R* = 0.41) (Extended Data Fig. 2c), which was much stronger for ASoC SNPs (*R* = 0.81) (Fig. 1c). Thus, our ASoC mapping in NGN2-Glut reproducibly identified putative functional SZ risk SNPs that altered chromatin accessibility in neurons generated from two different methods.

**Fig. 2:**
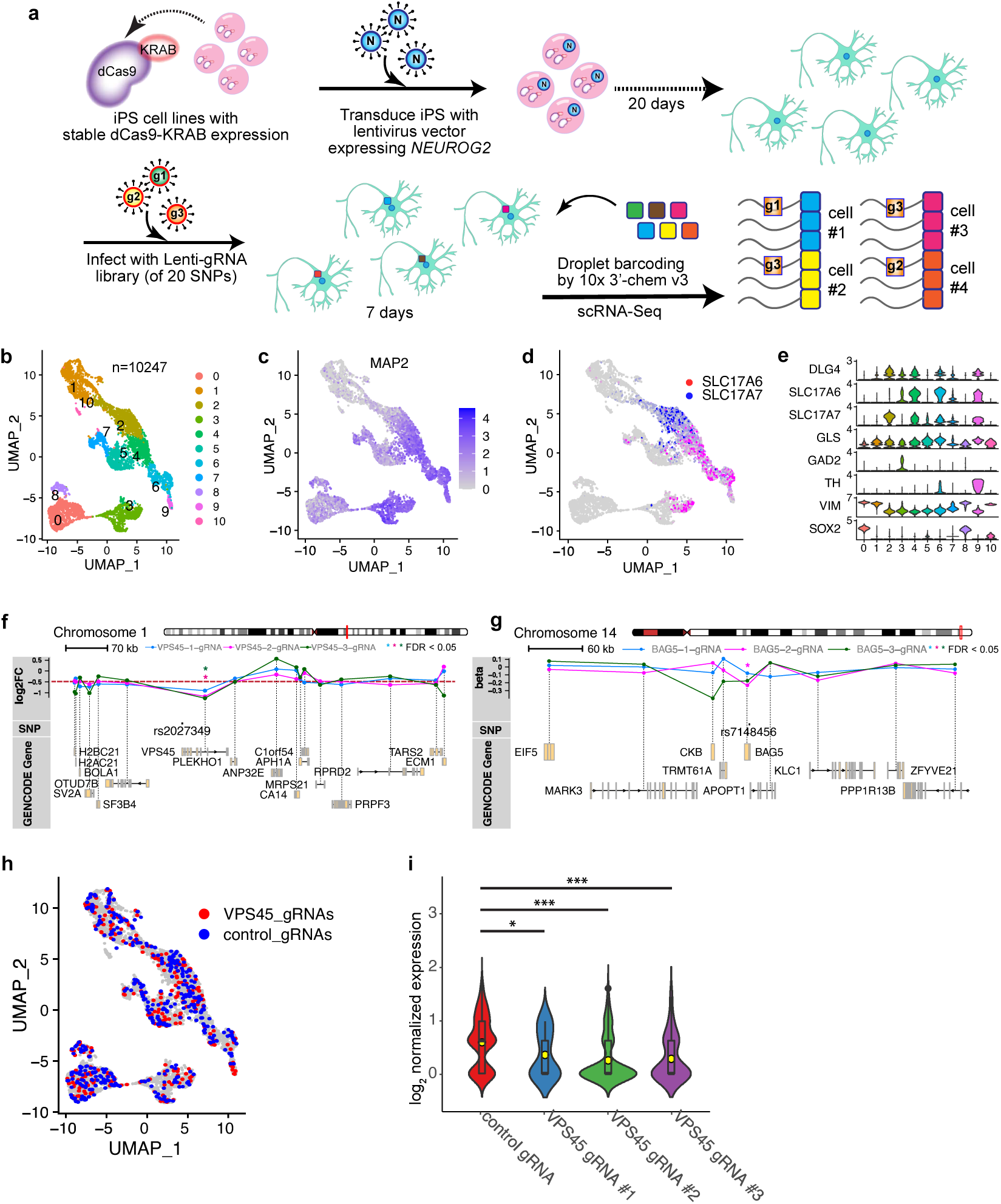
Multiplex CRISPRi combined with scRNA-seq in NGN2-Glut identifies regulatory sequences flanking SZ-associated ASoC SNPs. (a) Schematic of the modified CROP-seq approach to screen *cis*-targets of the 20 SNP sites in NGN2-Glut. (b) UMAP plot showing the 11 clusters of the 10,247 cells used in the scRNA-seq studies. (c) UMAP plot showing the normalized relative expression level of *MAP2* distribution. (d) UMAP plot showing the pattern of expression on glutamatergic neuron markers *SLC17A7* (vGlut1) and *SLC17A6* (vGlut2). (e) Violin plot set showing a set of different neuron-specific markers, including post-synaptic marker (*DLG4*), glutamatergic neurons (*SLC17A6*, *SLC17A7*, *GLS*), GABAergic neurons (*GAD2*), dopaminergic neurons (*TH*), and NPC-like cells with proliferative potential (*VIM*, *SOX2*). (f) Gene track-expression plot showing the effects of two CROP-seq gRNAs targeting rs2027349 (*VPS45* site) on the expression level change of their target and proximal genes. *: *FDR* < 0.05. (g) Same as (f), with the difference being three CROP-seq gRNAs targeting rs7148456 (*BAG5* site). (h) UMAP plot showing all cells assigned to either one of the three *VPS45* CROP-seq gRNAs (red) or one of the five control gRNAs (targeting to GFP; blue), note the homogenous distribution of *VPS45* and control gRNA-borne cells. (i) Quantification of the knockdown efficiency of *VPS45* CROP-seq gRNA (#1, #2, and #3 gRNAs) on the expression level of the *VPS45* gene. Kruskal-Wallis test (non-parametric) was used (shown are Dunn’s multiple comparisons adjusted *p*-values; ***: *P* < 0.001).

To nominate SZ risk SNPs for further functional study, we examined loci with GWS SZ-risk SNPs that showed ASoC in both datasets. Consistent with our previous SZ ASoC data in iN-Glut ^13^, rs2027349 at the *VPS45* locus remained the most significant ASoC in NGN2-Glut (Fig. 1d), followed by the ASoC SNP rs12895055 at the *BCL11B* locus (Supplementary Table 3, Extended Data Fig. 2d). rs2027349 at the *VPS45* locus is in very strong LD with the SZ GWAS index SNP rs12138231 (*R*^2^ = 0.94) and was fine-mapped as the only likely causal SNP at this locus in our recent study ^13^. The other two replicated ASoC SNPs also showing GWS association with SZ were at the *PBRM1*/*GNL3* (rs10933) and *BAG5* (rs7148456) loci (Supplementary Table 3, Extended Data Fig. 2d). These four SZ-associated ASoC SNPs were thus prioritized for functional testing for regulatory potential.

### Multiplex CRISPRi in NGN2-Glut confirmed the regulatory function of the ASoC sequence at the *VPS45* locus

We have previously performed multiplex CRISPRi combined with single-cell RNA-seq (scRNA-seq) (*i.e.*, a CROP-seq approach) in NPCs to screen regulatory sequences flanking ASoC SNPs ^13, 28^. Here, applying a similar approach in NGN2-Glut, we examined whether the OCR sequences that flank the four prioritized SZ-associated ASoC SNPs at *VPS45* (rs2027349), *BCL11B* (rs12985055), *PBRM1*/*GNL3* (rs10933), and *BAG5* (rs7148456) loci have regulatory potential, using the same set of guide RNAs (gRNAs) from our earlier study ^13^. NGN2-Glut stably expressing dCas9 fused to Krüppel associated box (KRAB) transcriptional repressor were transduced by lenti-gRNAs at low titer to maximize the chance that any neuron was infected by a single gRNA. The gene expression changes perturbed by each gRNA were assayed by scRNA-seq (Fig. 2a). NGN2-Glut appeared to be heterogenous, likely representing different maturity and possible subtypes of excitatory neurons (Fig. 2b-e, Extended Data Fig. 3a). To maximize the study power, we included all the MAP2+/SOX2-neurons of the Glut+ clusters (Fig. 2b-e; *N* = 8,921) in differential gene expression analysis, of which 4,057 neurons infected by a single gRNA (Extended Figure 3b and 3c).

**Fig. 3:**
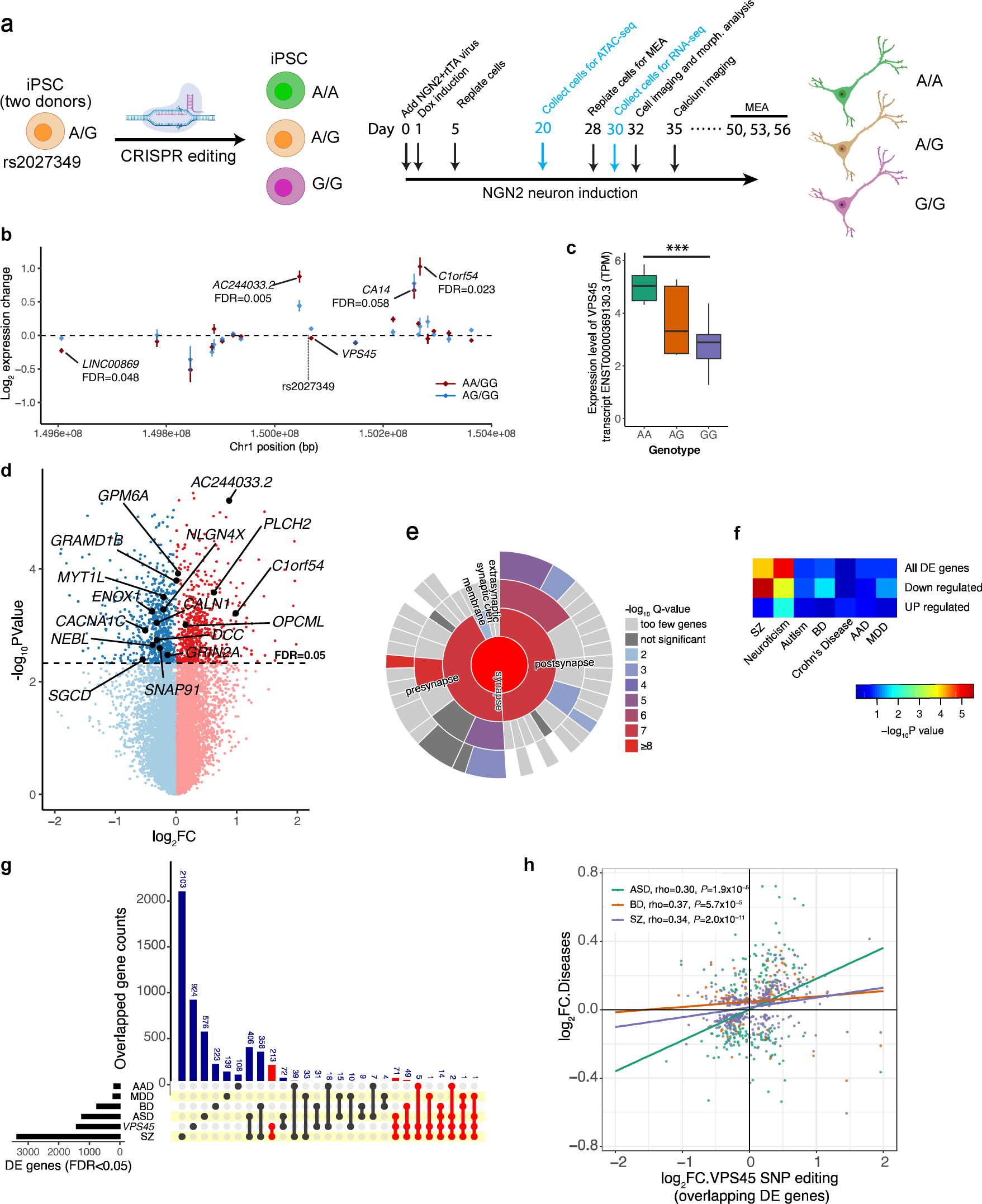
CRISPR/Cas9-editing of rs2027349 in NGN2-Glut results in local expression changes of multiple genes and transcriptome-wide disease-relevant expression changes. (a) Schematics of CRISPR/Cas9-editing of rs2027349 and neuron differentiation as well as a timeline of main experiments. (b) Different rs2027349 genotypes and their *cis*-effects on total RNA abundances of genes within a 500 kb window of rs2027349. The reference genotype (G/G) was used as the baseline. (c) A shift of G to A on rs2027349 leads to a significant increase of expression on *VPS45* transcript ENST00000369130.3. Kruskal-Wallis (non-parametric) test. ***: *P* < 0.001. (d) Volcano plot showing differential expression of the expressed genes (n = 14,999) in NGN2-Glut after rs2027349 editing. Highlighted are some DE genes (FDR < 0.05) that are also PGC3 SZ genes. The *P*-values were derived from a combinational analysis of all three genotypes (A/A, A/G, and G/G). -Log_2_FC was calculated as expression fold changes between A/A vs G/G neurons. (e) Sunburst plot showing strong enrichment of synapse-related ontology terms among the downregulated DE gene in A/A neurons as generated by SynGO ^71^. (f) MAGMA gene-set GWAS association enrichment analysis of DE genes for 6 neuropsychiatric disorders/traits and one unrelated disease (Crohn’s disease), showing high enrichment in SZ and neuroticism. (g) Upset plot showing the overlap of the DE genes from rs2027349 editing (A/A vs G/G neurons) against lists of DE genes associated with five neuropsychiatric disorders in postmortem brains. Note the high overlapping number (213) between *VPS45* and SZ sets. (h) XY-scatter plot showing the correlation of the log_2_ fold change of DE genes from rs2027349 (*VPS45* SNP) editing (A/A vs G/G neurons) against the overlapping DE genes of three major neuropsychiatric disorders (from g). Shown are corresponding ρ and *p-*values.

To identify the regulatory SNP site and its *cis*-target, we focused on genes within ±500kb from a targeting SNP/gRNA. We tested for differential expression (DE) of *cis*-genes between gRNA-targeted vs. non-targeted neurons using Student’s *t*-test, with *P*-values adjusted with the empirical null distribution obtained from permutation of cell labels (Supplementary Table 4). We found that sequences flanking three (rs2027349 near *VPS45*, rs10933 near *PBRM1*/*GNL3*, and rs7148456 near *BAG5*) of the four prioritized SZ-associated ASoC SNPs had *cis*-target genes with adjusted *P* < 0.05 (Fig. 2f-g; Extended Data Fig. 3d-e, Supplementary Table 4). All *cis*-targets with adjusted *P* < 0.05 showed reduced expression by respective gRNAs, consistent with the expected transcriptional repression by KRAB in CRISPRi ^29, 30^. *VPS45* near rs2027349 showed the most significant reduction of expression (Fig. 2h-i), which was consistent with rs2027349 exhibiting the strongest ASoC in NGN2-Glut among all the SZ risk SNPs (Figure 1b, Supplementary Table 3).

To assess which CRISPRi at the three loci (rs2027349, rs7148456, and rs10933, near *VPS45*, *PBRM1*/*GNL3*, and *BAG5*, respectively) resulted in global expression changes more biologically relevant to SZ, we first analyzed DE between gRNA-targeted vs. non-targeted neurons (Extended Data Fig. 4a), and then for each locus, we examined the up-/down-regulated genes for the enrichment of gene ontology (GO) terms (Extended Data Fig. 4b, Supplementary Table 5-7). We found that CRISPRi at the *VPS45* locus elicited the most biologically relevant transcriptomic changes, with the upregulated genes enriched for GO terms related to neural development and synaptic function (Extended Data Fig. 4b, Supplementary Table S5). For the *BAG5* locus, genes downregulated by CRISPRi were enriched for neuropeptide hormone activity and receptor-ligand activity, while for *PBRM1*/*GNL3* locus, neither up- nor downregulated genes showed enrichment of GO-terms relevant to neuronal function (Extended Data Fig. 4b, Supplementary Table 6-7). Our multiplex CRISPRi in NGN2-Glut thus confirmed the regulatory effects of the sequences flanking the ASoC SNPs at some SZ risk loci, in which CRISPRi at rs2027349 near *VPS45* showed the most pronounced effect.

### Precise allelic editing of the ASoC SNP at the *VPS45* locus reveals a complex gene regulation relevant to neurodevelopmental disorders

Given that rs2027349 near *VPS45* showed the strongest ASoC in NGN2-Glut (Fig. 1d), and that the DNA sequence flanking rs2027349 exhibited the most robust regulatory effect in CRISPRi (Fig. 2h-i), we next examined the functional effects of the precise allele editing of rs2027349 (Fig. 3a). We have previously performed CRISPR/Cas9 editing of two hiPSC lines heterozygous for rs2027349 from A/G to A/A and G/G ^13^. Here, with these CRISPR-edited isogenic lines of the three different genotypes (A/A, A/G, and G/G), we derived NGN2-Glut and performed transcriptomic DE analysis between different genotypes (Fig. 3a). We first examined the DE of *cis*-genes within a 500-kb interval of rs2027349 (Fig. 3b, Supplementary Table S8). We found *cis*-gene expression changes between A/A vs G/G neurons were correlated (Spearman’s Rho = 0.602) with those between A/G vs G/G (Fig. 3b and Extended Data Fig. 5a), confirming the consistency of the allelic effect in samples with different genotypes. Allele A of rs2027349 was associated with an increased expression of a *VPS45* transcript isoform (ENST00000369130.3) but not with total mRNAs of *VPS45* in NGN2-Glut (Fig. 3b-c, Extended Data Fig. 5b). Interestingly, we found allele A also significantly increased the expression of a lncRNA (*AC244033.2*) (FDR < 0.005) that is adjacent to *VPS45* but transcribed in the opposite direction, as well as a distal gene (*C1orf54*) that is ∼200 kb downstream of the edited SNP (FDR < 0.023) (Figure 3b, Supplementary Table 8). The increased expression of these *cis*-target genes of rs2027349 in A/A neurons was independently verified by quantitative polymerase chain reaction (qPCR) and was consistent in both donor lines (Extended Data Fig. 5c-f). These results suggest a complex *cis*-regulation of multiple genes (*VPS45*, *AC244033.2*, and *C1orf54*) by a single SZ risk SNP (rs2027349) at the *VPS45* locus.

*VPS45* has been suggested to play a role in vesicle-mediated protein trafficking and neurotransmitter release ^31^. While *AC244033.2* and *C1orf54* have no known function, *cis-* and *trans*-acting lncRNAs are known to be pervasive and may have profound roles in neurodevelopment ^32, 33^. We thus reasoned that the transcriptomic changes associated with the SZ risk allele A of rs2027349 might be relevant to neurodevelopment and synaptic function. Our DE analysis identified 687 upregulated genes and 731 downregulated genes (FDR < 0.05) in A/A neurons (vs G/G) (Fig. 3d). Gene set enrichment analysis (GSEA) using WebGestalt ^34^ showed that the upregulated genes were strongly enriched for GO terms related to neuron differentiation, neurogenesis, and nervous system development. In contrast, the downregulated genes were strongly enriched for GO terms such as synaptic signaling and synaptic membrane (Supplementary Table 9, Extended Data Fig. 6a-b). Because synaptic abnormalities have been implicated by the SZ GWAS (PGC Wave 3) ^1^, we further analyzed the DE genes by SynGO (synaptic gene ontology) ^35^ to confirm the relevance of transcriptomic changes to synaptic function. Consistent with the result from WebGestalt analysis, the downregulated genes in A/A neurons were significantly enriched for synaptic genes (pre- and post-synapse) (Fig. 3e, Supplementary Table 10). While the upregulated genes were not significantly enriched for synaptic genes (Extended Data Fig. 6c, Supplementary Table 11), some noteworthy upregulated genes are found in SynGO terms such as synaptic vesicle (*SLC17A7* and *SYN3*) and postsynaptic ribosome (Extended Data Fig. 6d). These results show that the SZ risk allele of rs2027349 affects global gene expression related to neurodevelopment and synaptic function.

We next assessed whether the transcriptional changes associated with the A allele of rs2027349 are relevant to neuropsychiatric disorders. We first examined the enrichment of DE genes for those associated with SZ. Of the 55 PGC3-prioritized single SZ risk genes ^1^ that were also expressed in NGN2-Glut, 10 were downregulated (*GRIN2A*, *MYT1L*, *NLGN4X*, *SNAP91*, *DCC*, *CALN1*, *NEBL*, *ENOX1*, *CACNA1C*, and *SGCD*), and 4 were upregulated (*PLCH2*, *OPCML*, *GPM6A*, and *GRAMD1B*) in A/A neurons (Fig. 3d, Supplementary Table 8), representing a 4.6-fold enrichment of SZ risk genes among downregulated genes (Fisher Exact Test *P* = 0.0002, two-tailed) and a 1.9-fold non-significant enrichment in upregulated genes (Extended Data Fig. 6e). Analysis of all the dysregulated genes within SZ loci in A/A neurons showed a significant enrichment of different SZ GWAS gene sets, and human post-synaptic density (PSD) gene sets often associated to SZ risk (Extended Data Fig. 6f) ^1, 26, 27, 37^. To account for potential confounding factors such as different gene sizes in GSEA and to orthogonally estimate the enrichment of DE genes for SZ GWAS risk, we performed MAGMA analysis using GWAS summary statistics ^38, 39^. We found that the downregulated genes in A/A neurons, but not the upregulated ones, showed strong enrichment of GWAS risk of SZ and, to a less extent, of neuroticism, bipolar disorder (BP), and major depressive disorder (MDD) (Fig. 3f). GWAS risk enrichment was not observed for autism spectrum disorders (ASD) likely due to the insufficient power associated with the small number of GWAS risk loci, or for other control disorders (*e.g.*, Crohn’s disease, CD) or for alcohol abuse disorder (AAD) (Fig. 3f) ^1, 4, 5, 40–43^. The results suggest that the transcriptomic changes associated with the SZ risk allele of rs2027349, especially the downregulated genes in A/A neurons, are relevant to SZ and other neuropsychiatric disorders.

To further ascertain the disease relevance of the transcriptional changes associated with the risk allele A of rs2027349 in isogenic NGN2-Glut, we examined whether the transcriptional changes in A/A neurons (vs. G/G) resemble the transcriptional differences in postmortem brain tissues of major psychiatric disorders. For overlapping DE genes (FDR < 0.05) between datasets (Fig. 3g), we found significant positive correlations between their expression changes in A/A neurons (*i.e.*, with rs2027349 edited at the *VPS45* locus) and in individuals of SZ (*ρ* = 0.34, *P* = 2 × 10^-11^), BP (ρ = 0.37, *P* =5.7 × 10^-5^), and ASD (ρ = 0.30, *P* =1.9 × 10^-5^) using the PsychENCODE datasets (Fig. 3g-h; Extended Data Fig. 6g) ^15^. The transcriptome-wide expression changes in A/A neurons and in individuals with SZ, BP, and ASD also exhibited significant correlations (ρ = 0.1-0.15, *P* =2.0 × 10^-37^ to 6.9 × 10^-68^; *N* = 13,816 genes) (Extended Data Fig. 6h). The expression changes in A/A neurons did not exhibit a significant correlation with transcriptional changes associated with MDD, or AAD (Extended Data Fig. 6g-h). Taken together, these results suggest that the ASoC SNP rs2027349 mediates a complex *cis*-gene regulation in NGN2-Glut that likely involves multiple genes (*VPS45, AC244033.2*, *and C1orf54*) at the *VPS45* SZ risk locus, subsequently leading to transcriptional changes relevant to neurodevelopment, synaptic function, and major neuropsychiatric disorders.

### Editing rs2027349 at the *VPS45* locus alters neurodevelopmental phenotypes in NGN2-Glut neurons

The effects of common SZ GWAS risk SNPs on human neuronal phenotypes have not been well established. Given the transcriptional effects of rs2027349-editing on synaptic and neurodevelopmental genes (Fig. 3), we examined whether rs2027349-editing alters neurodevelopmental phenotypes in NGN2-Glut neurons. Day-32 neurons differentiated from the isogenic SNP-edited lines (A/A, A/G, and G/G) were transfected sparsely with GFP to visualize the neuronal morphology (Fig. 3a), followed by Sholl analysis to examine genotype-specific effects on dendritic complexity (Fig. 4a). Two-way ANOVA (genotype × distance) showed that genotype had a significant effect on the number of intersections (dendritic branching) (*p* < 0.0001) (Fig. 4b). The number of intersections was significantly different at 80 to 160 μm from the soma, with G/G showing a reduced dendritic branching (Fig. 4a and 4b). A/A and G/G neurons were most different at 100μm from the soma (*P* < 0.0001, Šídák post hoc test).

**Fig.4:**
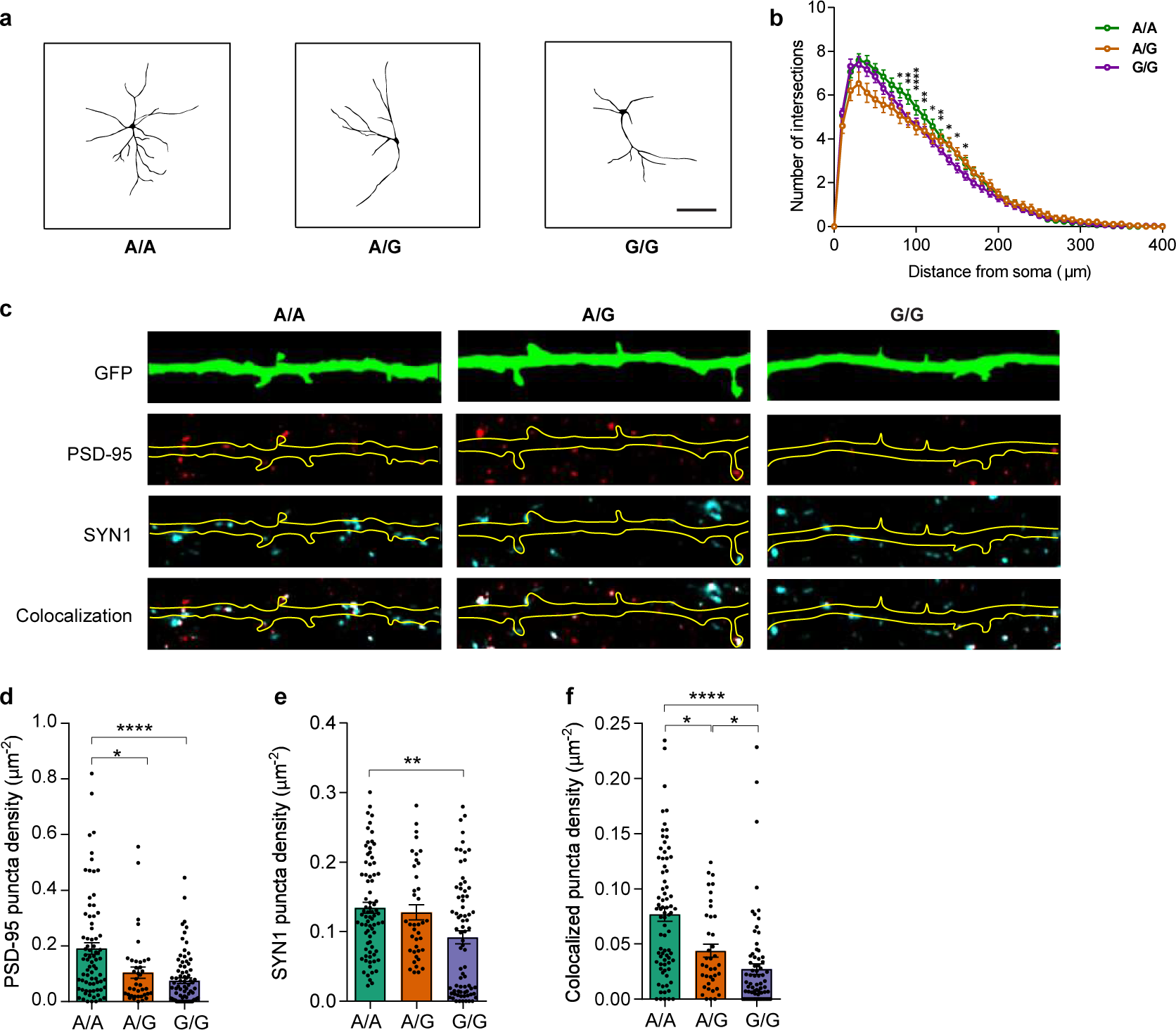
CRISPR/Cas9-editing of rs2027349 at *VPS45* locus alters neuron development in NGN2-Glut. (a) Representative traces of dendrites from rs2027349 A/A, A/G, and G/G neurons. Scale bar: 100 µm. (b) Sholl analysis of rs2027349 A/A (n=97), A/G (47), and G/G (n=101) neurons, showing that the A/A genotype is associated with increased dendritic complexity. Data points showing significant difference of distances between the A/A and G/G genotypes are marked on the graph. A two-way repeated measures ANOVA with Šídák multiple test correction was performed to determine significance. Data from 3 independent experiments were analyzed. (c) Representative immunofluorescence images of GFP-filled dendrites from NGN2-Glut neurons and synaptic puncta labelled with post-synaptic density 95 (PSD-95) and Synapsin I (SYN1). (d) Puncta density in neurons of different genotypes as measured by PSD-95 immunofluorescence. (e) Puncta density in neurons of different genotypes as measured by SYN1 immunofluorescence (f) Puncta density in neurons of different genotypes as measured by the colocalization of PSD95+/SYN1+, showing the A/A risk allele is associated with a higher density of excitatory synapses. Each dot represents synaptic density from one neuron (A/A, n=77; G/G, n=75; A/G, n=39). Data are from 2 independent experiments and were analyzed by a Kruskal-Wallis (non-parametric) test. *: *P* < 0.05; **: *P* < 0.01; ****: *P* < 0.0001.

We next examined the effect of rs2027349 editing on dendritic protrusions, which often host excitatory synapses (Extended Data Fig. 7a-b). We did not observe a significant difference in protrusion density between genotypes (Extended Fig. 7c-d). We then examined pre- and post-synaptic puncta on dendrites of NGN2-Glut neurons using immunofluorescence (Fig. 4c). We found that the density and area of both pre-synaptic (Synapsin 1; SYN1) and post-synaptic (PSD-95) puncta in A/A neurons were significantly higher than in G/G neurons (Fig. 4d-e; Extended Data Fig. 7e-g). The differences between A/A and G/G neurons were most robust for the colocalized SYN1^+^/PSD-95^+^ puncta density, labeling *bone fide* synapses. A/G neurons displayed an intermediate phenotype, significantly different from both A/A and G/G neurons in colocalized puncta density (Fig. 4f, Extended Data Fig. 7e). Thus, consistent with the transcriptional effects of the rs2027349 editing, these results suggest that the ASoC SNP rs2027349 affects dendritic complexity and synapse development in NGN2-Glut.

### Editing rs2027349 at the *VPS45* locus affects synaptic function in NGN2 neurons

To ascertain whether the neurodevelopmental changes in rs2027349-edited neurons were accompanied by altered neuronal function, we assessed neuronal network activity in isogenic NGN2-Glut carrying different genotypes of rs2027349. We used multielectrode arrays (MEAs) to evaluate neuronal population dynamics and calcium (Ca^2+^) imaging to assess single-cell activity (Fig. 3a, Fig. 5a-b). Using MEAs, we found that isogenic neurons exhibited differences for mean firing rate and the number of bursts between days 41 to 74 (after neural induction), a time when neurons showed the most active firing (Fig. 5c-d; Extended Data Fig. 8a). At days 50, 53, and 56, the mean firing rates were about 50% higher (*p* values = 0.004-0.006, Student’s *t*-test, two-tailed), and the average number of bursts per 10 min were about 65% more (*p* values = 7.8 × 10^-5^ to 4.0 × 10^-6^, Student’s *t*-test, two-tailed) in A/A compared to G/G neurons, with A/G neurons displaying an intermediate phenotype (Fig. 5c-d). Network burst frequency was only found to be different at day 50, suggesting the impact may be restricted to specific neuronal developmental stages. The synchrony index was not different between genotypes, despite observing a general increase in synchrony during later stages of network development (Extended Data Fig. 8a).

**Fig. 5.**
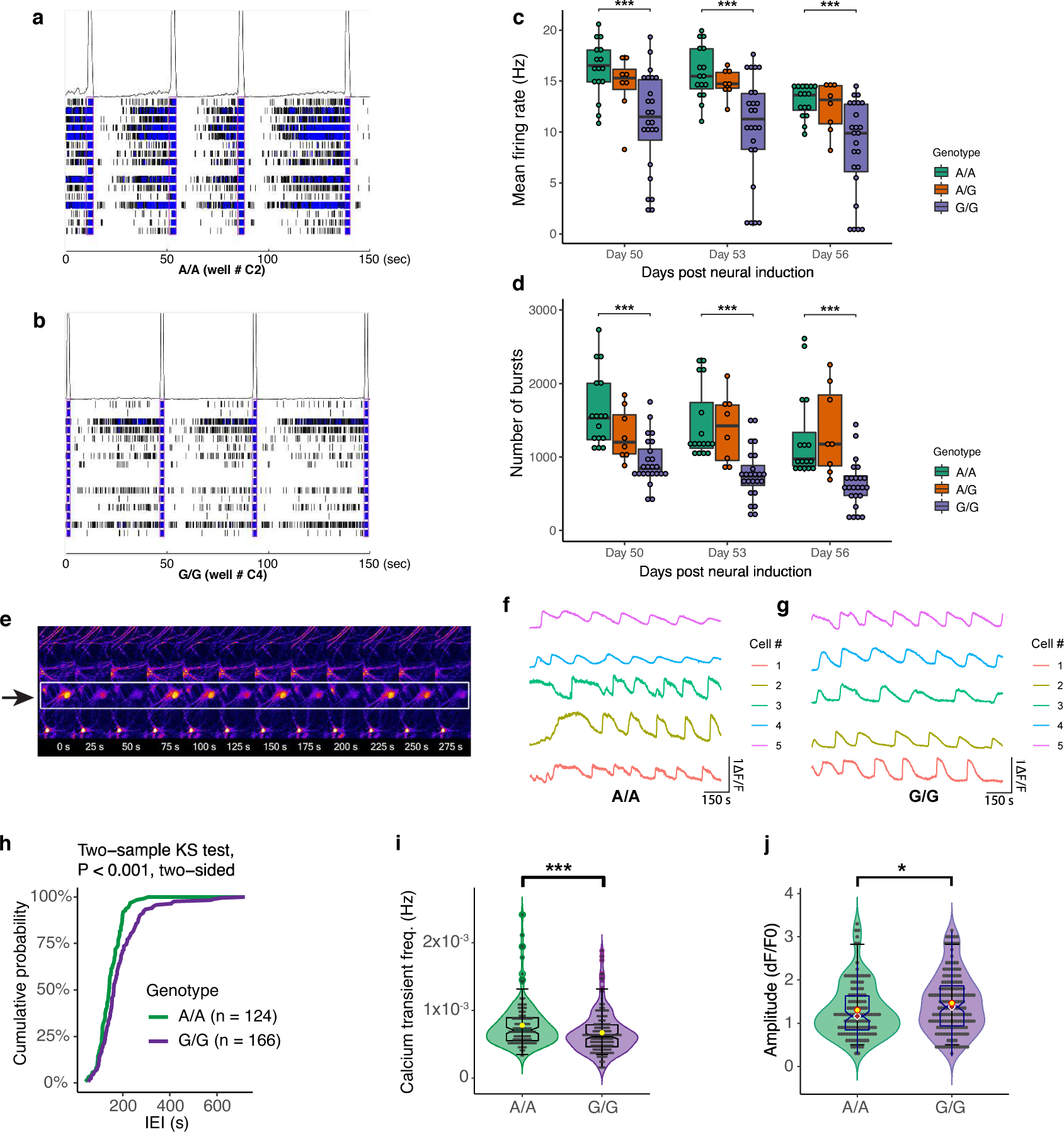
CRISPR/Cas9-editing of rs2027349 alters neural network and electrophysiological activity of NGN2-Glut. (a-b) Raster plots of neuronal firing events assayed by multi-electrode array (MEA) with 16 electrodes per well. (a) Well C2 from genotype A/A. (b) Well C4 from genotype G/G. (c-d) Statistics of mean firing rate and burst numbers at day 50, 53, and 56 in NGN2-Glut with different genotypes of rs2027349. Each dot represents data collected from one MEA well from two independent experiments. Kruskal-Wallis (non-parametric) test. ***: *P* < 0.001. (c) Mean firing rate, showing a significant increase in firing rate in A/A neurons. (d) The number of bursts, showing a significant increase in burst numbers in A/A neurons. (e) Representative two-photon pseudo-color Ca^2+^ imaging time-series images showing a neuron and its firing activities in a 275-second (275 s) span. (f-g) Representative neuron firing signals from five cells of each genotype are indicated in (f) A/A and (g) G/G. (h-j) Ca^2+^ transient frequency and amplitudes in A/A (*n* = 124) and G/G (n=126) neurons from two independent experiments. (h) Cumulative probability plot of Ca^2+^ transient inter-event intervals (IEI) in A/A and G/G neurons, two-sample KS test, two-sided. *P* < 0.001 (i) Violin and box plot showing the distribution of Ca^2+^ transient frequency in A/A and G/G neurons. Yellow dot: mean value. Student’s *t*-test (non-parametric, two-tailed), *P* < 0.001. (j) Violin and box plot showing the distribution of Amplitude (dF/F0) in A/A and G/G neurons. Yellow dot: mean value. Student’s *t*-test (non-parametric, two-tailed), *P* < 0.001.

We next assessed the effect of SNP editing on individual neuron activity by Ca^2+^ imaging. We used a cell-membrane permeable Ca^2+^ indicator (Fluo-4 AM) to measure intracellular Ca^2+^ dynamics in A/A and G/G neurons at day 35 after neuron induction (Fig. 3a, 5e-g). We found that the neuronal Ca^2+^ transient (firing) frequency in A/A neurons was significantly higher than in G/G neurons (*P* < 0.001, Student’s t-test, two-tailed) with shorter inter-event intervals in A/A neurons (*P* < 0.001, two-sample KS test) (Fig. 5h-i). There was also a significant (*P* = 0.042) reduction in the amplitude of Ca^2+^ transients in A/A neurons (Fig. 5j). We found that Ca^2+^ transients were strongly correlated (synchronization index ranges from 0.7 to 0.9) between different neuronal cultures for each genotype; however, no significant difference in synchrony index was found between A/A and G/G neurons (Fig. 5f-g, Extended Data Fig. 8b-c). The observed higher frequency of Ca^2+^ transients in A/A neurons was consistent with the increased population neuronal firing rate from MEA in A/A neurons (Fig. 5a-d). Together, these results suggest that SZ risk allele of rs2027349 is associated with hyperactivity of NGN2-neurons, which is congruent with some previously reported neuronal hyperactivity associated with SZ risk alleles ^12, 14, 44, 45^.

### All three genes, *VPS45, AC244033.2*, and *C1orf54,* at the *VPS45* locus contribute to the altered neural phenotypes in a non-additive fashion

Given that the ASoC SNP rs2027349 can *cis*-regulate multiple genes (*VPS45, AC244033.2*, *and C1orf54*) (Fig. 3b-c) and alter neuronal phenotypes (Figs. 4-5), we attempted to determine which gene(s) at this locus likely contributed to the SZ risk allele-associated neuronal phenotypes. To this end, we carried out individual gene knockdown (KD) by stably expressing lentivirus-based short-hairpin RNAs (shRNAs) targeting *VPS45, AC244033.2, or C1orf54* (with shRNAs targeting *EGFP* as a control) in CRISPR-edited A/A hiPSCs, which were then differentiated into NGN2-Glut for assessing whether the risk allele-associated cellular phenotypes could be reversed. We first confirmed the expected reduction of *VPS45, AC244033.2, or C1orf54* expression in NGN2-Glut for each individual gene KD (Fig. 6a-c, Extended Data Fig. 9a, and Supplementary Table 12). We then assessed the effect of KD in A/A neurons to mimic the phenotypes observed in G/G neurons. We found that shRNA of *C1orf54 (*shRNA*_C1orf54),* and to a less extent shRNA of *AC244033.2 (*shRNA*_lncRNA)* reversed dendritic complexity in A/A neurons towards the level of G/G neurons (Fig. 6d, Extended Data Fig. 9b). While we found that *VPS45* KD did not alter dendritic complexity (Fig. 6d; Extended Data Fig. 9b), our Ca^2+^ imaging showed that KD of *VPS45*, as well as *lncRNA* and *C1orf54*, all partially reversed the higher Ca^2+^ transient frequency of A/A neurons towards the level of G/G neurons (Fig. 6e). Together, these results suggest that *VPS45, AC244033.2*, *and C1orf54* all contribute to the phenotypic changes in rs2027349-edited A/A neurons.

**Fig. 6:**
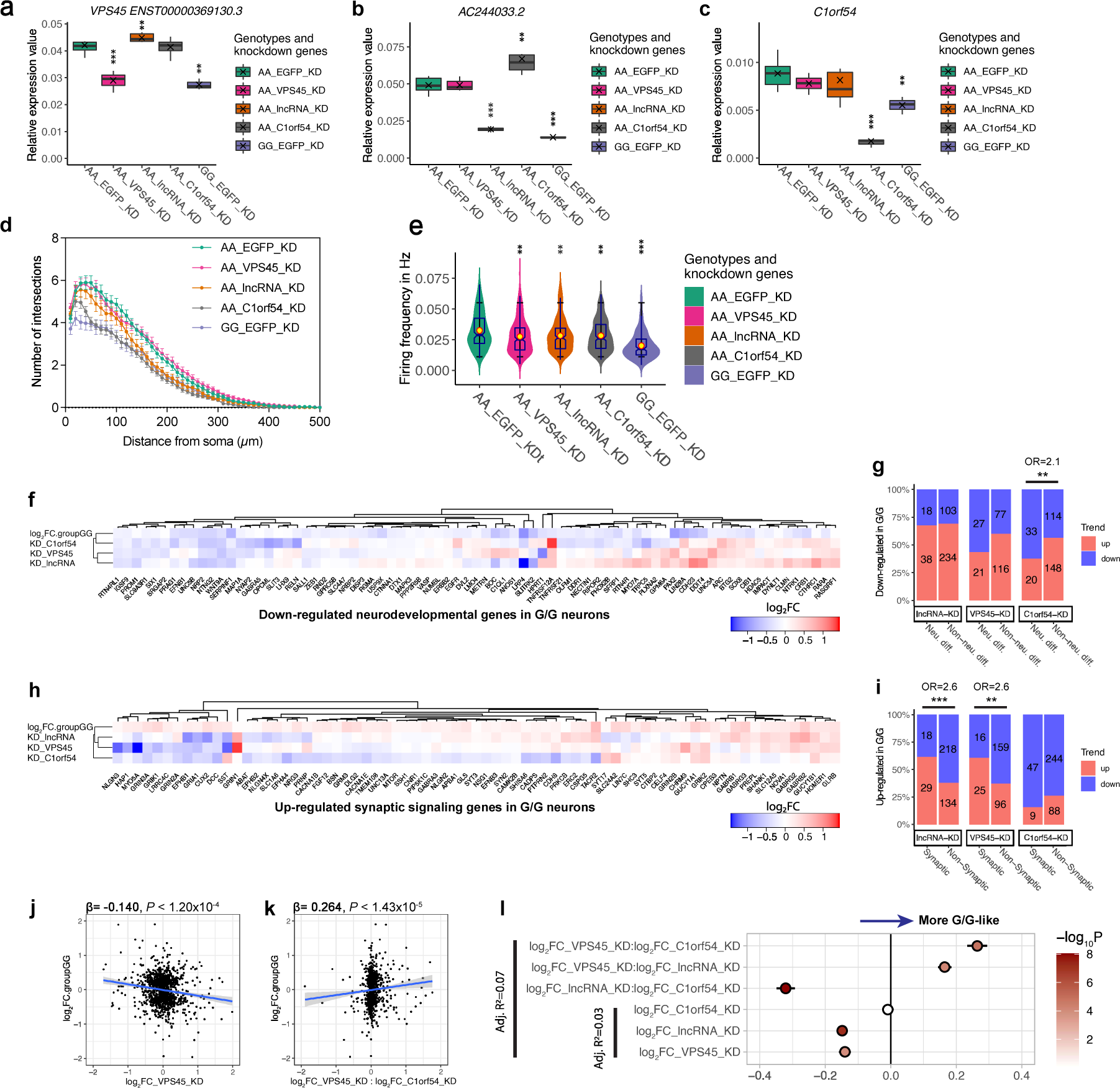
*VPS45*, *AC244033.2 (lncRNA)*, and *C1orf54* interactively contribute to the altered neural phenotypes in NGN2-Glut at the rs2027349 locus. (a-c) Box-whisker plots showing shRNA-mediated KD of each gene in A/A neurons reversed gene expression to a level more similar to that G/G neurons. The expression was measured by qPCR with GAPDH as the endogenous control for normalization. In each subpanel, the comparison was made using AA_EGFP serving as control. Saltire mark: mean value. Kruskal-Wallis test (non-parametric). *N* = 3-6 biological replicates in each experiment. (a) Knockdown of *VPS45* gene recapitulated *VPS45* transcript ENST00000369130.3 expression pattern in G/G neurons. (b) Knockdown of *AC244033.2* recapitulated its expression pattern in rs2027349 G/G neurons. (c) Knockdown of *C1orf54* recapitulated its expression pattern in rs2027349 G/G neurons. (d) Sholl analysis revealed knockdown of *AC244033.2* or C1orf54 in A/A neurons partially reversing its dendritic complexity to the level of G/G neurons, see Extended Data Fig. 9b for statistics. (e) Knockdown of *AC244033.2* or *C1orf54* in A/A neurons reduced firing frequency and partially reversed it to the level of G/G neurons. Kruskal-Wallis (non-parametric) test with Dunn’s multiple comparisons and adjusted *P*-values. *N* = 181, 245, 229, 276, and 152 neurons from two independent experiments. **: *P* < 0.01. (f) Hierarchical clustering (nearest distance method) of log_2_FC for neurodevelopmental genes downregulated in rs2027349-edited G/G neurons and their expression changes in each shRNA KD in A/A neurons. (g) The enrichment of the up-/down-regulated genes in each shRNA KD in A/A neurons among the neurodevelopmental (*i.e.*, neuron differentiation related) genes downregulated in rs2027349-edited G/G neurons. (h) Same as (f), using synaptic genes upregulated in G/G neurons. (i) Same as (g), examining the enrichment among synaptic genes upregulated in G/G neurons. In (g) and (i), Fisher’s exact test (two-sided) was used to estimate the enrichment; OR: odds ratio, ***: *P* < 0.001. (j) Scatter plot showing the correlation and linear regression result between the log_2_FC of DE genes in rs2027349-edited G/G neurons (G/G vs A/A) and the log_2_FC in *VPS45* shRNA KD conditions in A/A neurons. (k) Similar to (j), using a linear regression model introducing the interaction of log_2_FC from *VPS45* KD and *C1orf54* KD against the log_2_FC of DE genes in rs2027349-edited G/G neurons. (l) Dot and whisker plot showing the summary of correlations from linear regression modeling fitting, indicating a non-additive effect of each gene KD in explaining the transcriptional effects of rs2027349-editing. Note that the interaction terms of shRNA KD (*VPS45*:*C1orf54* and *VPS45*:*AC244033.2*) demonstrated a strong positive coefficient as in the rs2027349-editing effect in G/G neurons. Whisker: ±95% CI; fill: *P*-value.

To further understand how each gene (*VPS45, AC244033.2*, *or C1orf54)* contributes to the neural phenotypic effects, we examined their impacts at the molecular level. We performed RNA-seq for each KD condition in two cell lines (Extended Data Fig. 9c, d) and compared the list of DE genes of each KD condition with DE genes in rs2027349-edited G/G (vs. A/A) neurons (Supplementary Tables 12, 13). Since the number of the DE genes (FDR < 0.05) for each KD condition was large, with 28%, 49%, and 44% of the expressed genes for KD of *VPS45, AC244033.2*, and *C1orf54,* respectively (Extended Data Fig. 9d), we focused our analyses on a subset of DE genes most relevant to our observed neuronal phenotypic changes by rs2027349 editing. These selected genes belong to the enriched GO-terms (from rs2027349-editing) related to neuron development and neuron differentiation (downregulated in G/G) or those related to synaptic signaling (upregulated in G/G) (Fig. 6f-i, Supplementary Tables S9, S13). We then asked whether each subset of those DE genes was enriched for any shRNA KD-associated DE genes with the directional expression change consistent with the rs2027349-editing effect (Fig. 6f-i). We found that neuronal differentiation genes with reduced expression in SNP-edited G/G neurons were significantly enriched for genes downregulated by shRNA KD of *C1orf54* (*OR* = 2.1, *P* = 0.009; Fisher’s exact test) but not with *AC244033* or *VPS45* (Fig. 6f-g, Table S13). For synaptic signaling genes with increased expression in SNP-edited G/G neurons, we observed an excess of genes upregulated by shRNA KD of *VPS45* (OR = 2.6, *P* = 0.004; Fisher’s exact test) and *AC244033.2* (*OR* = 2.6, *P* = 0.001; Fisher’s exact test) but not with *C1orf54* (Fig. 6h-i, Table S13). These results provide mechanistic support for the observation that all three genes contribute to the phenotypic changes in rs2027349-edited neurons, likely through influencing the expression of different sets of biologically relevant genes.

We next tested if the transcriptional effects of rs2027349-editing could be explained by the effects of individual KD (*VPS45, AC244033.2*, or *C1orf54)* in NGN2-Glut. We first fit a linear regression model where we regressed expression fold-changes from the SNP editing, as a response variable, against expression changes from KD of three individual genes. For all the DE genes from the SNP editing (G/G vs A/A) (Supplementary Table 13), our model fitting explained 3% of the SNP editing effects across genes (adjusted *R*^2^ = 0.03, *P* < 4.3 x 10^-9^), with an unexpected negative correlation (Fig. 6j, Extended Data Fig. 9e). However, when we introduce the interaction terms between the expression effects of individual KD, we observed a marked improvement of model performance, explaining 7% of the variation of the SNP editing effects (adjusted *R*^2^ = 0.07, *P* < 2.2 x 10^-16^). The gene × gene interaction terms were highly significant, with *VPS45*_KD × *C1orf54*_KD (b = 0.26, *P* = 1.4 × 10^-5^) and *VPS45_*KD × *lncRNA*_KD (b = 0.16, *P* = 6.7 × 10^-5^) positively correlated with the SNP editing effect (Fig. 6k, l, Extended Data Fig. 9f). The highly significant interaction terms of different shRNA KD were consistent with our further GO-term enrichment analysis of DE genes from each shRNA KD, where we found that overlapping genes between different KD conditions showed stronger enrichment for biologically relevant GO-terms, *i.e.*, relating to neurodevelopment and synaptic signaling (Extended Data Fig. 10; Supplementary Table 14). These results suggest that *VPS45, AC244033.2*, and *C1orf54* contribute to the phenotypic changes in rs2027349-edited neurons in a non-additive fashion.

Finally, to mechanistically understand the *cis*-SNP editing effect on all three genes, we examined the existing brain chromatin interactions (*i.e.*, capture Hi-C) ^46^ and our current ATAC-seq data (A/A vs G/G neurons) around the edited rs2027349 site. We found that the rs2027349 site had long-range chromatin contacts with promoter regions of many genes, including *VPS45/AC244033.2* and *C1orf54* (hippocampal neurons ^46, 47^; Extended Data Fig. 11a), which supports the *cis*-regulatory effect of rs2027349-editing on the distal *C1orf54* in NGN2-Glut. For local chromatin accessibility in NGN2-Glut, we found that rs2027349 editing altered the enhancer/promoter OCR peaks of both *VPS45/AC244033.2* and *C1orf54* loci, with A/A neurons showing higher regional OCR peaks than in G/G neurons (Extended Data Fig. 11b). The higher regional OCR peaks at *VPS45/AC244033.2* and *C1orf54* loci were consistent with the increased expression of all three genes by SNP editing in A/A neurons (Fig. 3b-c). Taken together, the results from the neuronal phenotypic reversal by individual gene KD and from the integrative transcriptomic/epigenomics analyses suggest an intriguing gene regulation model at a single SZ GWAS risk rs2027349 locus, where all three genes (*VPS45, AC244033.2*, and *C1orf54*) contribute to the neuronal phenotypes in a non-additive fashion.

## Discussion

Functional interpretation of noncoding GWAS risk variants is a major challenge in genetics. Combining analyses of allelic chromatin accessibility, CRISPRi screening, and precise SNP editing in NGN2-Glut, we prioritized regulatory SZ risk variants affecting neural chromatin accessibility and gene expression. For the functional SZ risk variant at the *VPS45* locus, we tied the SNP function to neural phenotypic changes. More importantly, we demonstrated a complex gene regulation paradigm where a single SZ risk locus mediates expression of both proximal and distal genes, which synergistically confer non-additive effects on disease-relevant cellular phenotypes.

Long-range gene regulation for neuropsychiatric genetic risk variants has been supported by brain promoter-capture Hi-C study ^46^. However, such postulated long-range *cis*-regulatory effects involving multiple genes and its relevance to cellular phenotypes has not been well established. Here, we showed that a single GWAS risk SNP (rs2027349) at the *VPS45* locus exhibited strongest ASoC and affected the expression of both its adjacent genes (*VPS45* and *AC244033.2*) and a distal gene *C1orf54 (*∼200 kb away) (Fig. 3b-c). The long-range effect of rs2027349 on *C1orf54* in NGNs-iNs was not observed in NPCs ^13^, supporting the notion of cellular state-specific regulation of gene expression ^13, 18, 46^. We also found that all three genes *cis*-regulated by rs2027349 contribute to the cellular phenotypic changes (Fig. 6). Interestingly, the transcriptional effect of rs2027349-editing can be better explained by gene × gene interactions (*VPS45* × *C1orf54* and *VPS45_*KD × *lncRNA*_KD) than by individual genes (Fig. 6g). The observed non-additive effects of multiple genes at a single GWAS risk locus is analogous to recently reported synergistic transcriptomic effect of SZ risk SNPs from different loci ^12^. Collectively, our results provide empirical support for a complex gene regulation with compound effects from multiple genes at a single SZ GWAS risk locus.

Disease-relevant cellular phenotypes for neuropsychiatric disorders, especially for common GWAS risk variants, remain largely undefined. Here, with isogenic rs2027349-edited NGN2-Glut, we tied the SZ risk allele of rs2027349 with increased dendritic complexity, synaptic puncta maturation, and neuronal activity (Figs. 4-5). The increased neuronal function by the risk allele of rs2027349, although seemingly inconsistent with the documented reduction of dendritic complexities and synaptic function for SZ ^48, 49^, is similar to the effect of a SZ GWAS risk variant at the *MIR137* locus ^14^, as well as the effects of rare SZ risk variants with high penetrance (*e.g.*, 16p11 duplication or loss-of-function of *SHANK3*) ^44, 45^. Together, with other studies that reported both neural hyperfunction ^12–14, 44, 45^ and hypofunction ^48, 49^ in SZ, our results provide further support for a neuronal homeostatic model of neuropsychiatric disorders where either excess or inadequate synaptic signaling output may contribute to pathophysiology ^50, 51^.

Although our observed effect of rs2027349-editing on neuronal phenotypes and the involvement of all three *cis*-target genes were supported by transcriptomic analyses, these results still need to be interpreted cautiously. This is because DE analysis, even for isogenic hiPSC lines, can be confounded by clonal variations ^56^, which is a general challenge for the field. However, such limitations are somewhat mitigated by our use of multiple lines/clones of all three rs2027349 genotypes (A/A, A/G and G/G). The confidence on our results was further enhanced by the observed congruence of GO analysis of the DE gene sets with the neuronal phenotypic alterations in rs2027349-edited neurons, as well as by the mechanistic support from the direct chromatin contacts and co-regulation of chromatin accessibility between rs2027349 and its three *cis*-target genes (Extended Data Fig. 11). Collectively, our study uncovers a complex gene regulation at a single SZ GWAS risk locus where an ASoC SNP alters chromatin accessibility of multiple putative risk genes that have non-additive cellular transcriptional and phenotypic effects in excitatory neurons, providing a novel mechanistic link between a noncoding SZ GWAS risk variant and disease-related phenotypes.

### Online Methods

(For more details on the experimental methods, please refer to the Supplementary Notes)

#### Selection and culture of hiPSC lines

The 20 hiPSC lines used for generating NGN2-Glut neurons for ATAC-seq were the same lines as used in our previous study ^13^. These hiPSCs were derived from cryopreserved lymphocytes (CPLs) using the genome-integration-free Sendai virus method, and the purity was confirmed by both immunofluorescence (IF) and Pluritest (https://www.pluritest.org/) with RNA-Seq data. Lines CD0000011 and CD0000012, which are heterozygous (A/G) at SNP site rs2027349, were used for CRISPR/Cas9 editing into all three genotypes, 2-3 clones per line ^13^. The hiPSCs were maintained in mTeSR Plus media (STEMCELL) on Matrigel (Corning) coated 6-well plate with media refreshment every other day. Cells were passaged at 1:10-1:20 every 4-6 days using ReLeSR (STEMCELL) following the vendor’s protocol. The NorthShore University HealthSystem Institutional Review Board (IRB) approved the study.

#### Generation of NGN2-Glut neurons

NGN2-Glut neurons were prepared as described ^20^ with modifications. Briefly, on Day 0, 5 × 10^5^ hiPSCs were dissociated using Accutase (STEMCELL) and replated in 1.2 ml mTeSR Plus media containing 5 µM ROCK inhibitor, rtTA virus, and NGN2-puro or NGN2-Hygro virus on 6-well plates. On Day 1, media was refreshed with mTeSRPlus containing 2 µg/ml doxycycline. From days 2 to 4, media was refreshed daily with Neurobasal Medium supplemented with 1× Glutamax/B27, 2 µg/ml doxycycline, and 2 µg/ml puromycin or 200 µg/ml hygromycin. On day 5, cells were dissociated with Accutase, resuspended in Neurobasal Medium supplemented with 1× Glutamax/B27, 2 µg/ml doxycycline, and 10ng/ml BDNF/GDNF/NT-3, and subsequently replated at 2 × 10^5^/cm^2^ on 0.1% PEI/Matrigel coated 12-well plates. For morphological immunostaining, 5 × 10^4^ iNs were replated onto 12 mm coverslips (with 5 × 10^4^ rat astrocytes pre-cultured) in Neurobasal Medium supplemented with 1× Glutamax/B27, 2 µg/ml doxycycline, 10 ng/ml BDNF/GDNF/NT-3, and 1% FBS. From Day 6, media was refreshed every three days with a half volume change. Doxycycline was withdrawn from Day 14 onwards. Sample harvesting for different purposes was performed at different times (Fig. 3A). For cells cultured on coverslips, GFP transfection was performed at Day 30 using Lipofectamine 3000 (Thermofisher) at 1:2 DNA:reagent ratio with 1 µg plasmid per coverslip. On Day 32, coverslips were fixed and stained for morphological analysis.

#### qPCR analysis of gene expression levels

RNA was isolated using RNeasy plus kit (QIAGEN) following the vendor’s instructions and quantified by a NanoDrop-8000 spectrometer. RT was performed using the High-Capacity cDNA Reverse Transcription Kit (Thermofisher) following the vendor’s instructions. qPCR was performed on a Roche 480 II Lightcycler using gene-specific FAM-labelled TaqMan probes with *GAPDH*, *ABL1*, and *CASC3* serving as internal controls. Biological replicates (3 to 6) were included for all conditions, and three technical replicates were used during qPCR.

#### RNA-seq data generation and analysis

Total RNA was isolated using RNeasy plus kit (QIAGEN) following the vendor’s instructions and quantified by a NanoDrop-8000 spectrometer. The isolated RNA was subsequently sent to the sequencing provider (GENEWIZ) for further processing and sequencing analysis. The data was returned as 2×150 bp paired-end fastq files. Raw files were subsequently mapped to the human hg38 genome (GRCh38p7.13) and Gencode v35 using STAR ^57^ v2.7.0. Direct quantification using the built-in function of STAR and probabilistic quantification by Kallisto ^58^ were used for different purposes in separate experiments. DE gene analysis was performed using EdgeR ^59^ by fitting the dataset into a gene-wise negative binomial generalized linear model with a quasi-likelihood evaluation method, with cell lines and batch numbers introduced as blocking factors whenever possible. When computing PCA projections, the top 15,000 DE genes were used.

#### ATAC-seq data generation analysis

ATAC-seq samples were prepared as previously described ^13, 60^. Briefly, 75,000 viable cells were lysed and nuclei collected for Tn5 reaction per sample. Transposed DNA was purified by DNA Clean and Concentrator-5 Kit (Zymo), and the eluted DNA was subsequently shipped to the University of Minnesota Genomic Center for library preparation and ATAC-seq.

Raw reads were provided as 2×75 bp paired-end fastq files targeting 60 M depth per sample. Raw files were pre-trimmed and mapped to the human genome reference file, including decoy sequences (GRCh38p7.13/hg38, 1000 Genome Project) by bowtie2. Picard v2.23 was used to remove all PCR and optical duplicated reads from the BAM file. To further eliminate allelic bias towards reference alleles during the aligning step, we performed WASP calibration on the generated raw BAM files ^61^ to get a set of calibrated BAM files for subsequent peak detection and SNP calling. The BAM file from each sample was first sub-sampled to equal size (30 M) to eliminate sample size-induced bias, and the 20 sub-sampled BAM files were merged as the source file for peak calling. MACS2 ^25^ was used to generate peak files (narrowPeak format) with recommended settings at FDR = 0.05. GATK (version 4.1.8.1) was used for ASoC SNP calling, as recommended by the GATK Best Practices (software.broadinstitute.org/gatk/best-practices/)^62^. When detecting ASoC SNPs, we pooled SNPs for all the called heterozygous SNPs locations from the 20 samples as justified by the high concordance of allele-specific effects across individuals ^13^, and only biallelic SNP sites (GT: 0/1) with minimum read depth (DP) ≥ 20 and minimum reference or alternative allele count ≥ 2 were retained. The binomial *P*-values were calculated with the Benjamini & Hochberg correction at *P* < 0.05 to define significant ASoC SNPs. When comparing the changes of chromatin accessibility caused by genotypes across samples, read counts were scaled and normalized using the deepTools package `bamCoverage` function and re-scaled to reads per genomic content (RPGC) as the base unit ^63^. The PCA graph (Extended Data Fig. 1g) used the chromatin accessibility data from our previously published studies ^13^, as well as DNase I hypersensitive assay from the fetal brain (NIH Roadmap) ^64^ projected to our published cross-cell-type non-overlapping OCR peak interval (n = 666,614) to integrate our NGN2-Glut into the multi-type datasets. (See Supplementary Materials and Methods for all the parameters and details).

#### CROP-seq in NGN2-Glut neurons

Lentivirus particles were packaged in fresh HEK 293T culture (passaged ≤15 times) as previously described ^13^. Transfection was carried out using FuGENE HD (Promega) following the vendor’s instructions. The virus-containing media was aliquoted into 1.5 ml low protein binding tubes and stored at -80°C until use.

The hiPSC lines (CD000009 and CD000011) that stably express dCas9-KRAB-BSD were established as previously described ^13^. NGN2-Glut were prepared from hiPSCs stably expressing dCas9-KRAB-BSD using NGN2-hygro virus as described in the section “Generation of NGN2-Glut neurons”. On Day 20, neurons were replated and infected with CROPseq-Guide-Puro virus (vector carrying gRNA targeting genes of interest, MOI=0.2) as described in the original CROP-seq article ^28^. Puromycin selection started 48 hrs after transduction. 7 days after selection, cells were harvested for single-cell RNA-seq preparation.

10X Genomics Cell Ranger v4 was used to process raw sequencing data. Briefly, sequencing data were aligned to the human GRCh38/hg38 genome with spike-in gRNA sequences as artificial chromosomes (20 bp gRNA sequence and 250 bp downstream plasmid backbone per each gRNA, together with one additional full sequence of CROP-Seq-Guide-Puro construct also included as a decoy to capture vector-borne sequences). The filtered gene count matrix output from Cell Ranger was used in subsequent R analysis. The R package Seurat v 3.2 was used to extract information and assemble data tables from the output files to generate the digital gene expression matrix, which was constructed based on per-gene UMI count. A cell assigned with a unique RNA was defined as the UMI count of its dominant gRNA in the cell being at least three times more than the sum of UMI counts from all other gRNAs. Only MAP2+ cells with strong glutamatergic neuron marker genes (SLC17A6, SLC17A7) falling within clusters 2, 3, 4, 5, 6, and 7 were used (Fig. 2b). Dimension reduction, UMAP, clustering, and pseudo-bulk DE analysis were performed using package Seurat and EdgeR. Graphs were made in Seurat and ggplot2.

#### Immunostaining and microscopy

Characterization of hiPSCs and neurons was performed as described ^13^ with minor modifications. Briefly, fresh cells were fixed in 4% PFA at room temperature, washed, permeated by 1% Triton X-100 in PBS, and blocked with 3% BSA. Samples were incubated with primary antibody at 4°C overnight and secondary antibodies for 1 hr at room temperature. Samples were mounted using ProLong™ Diamond Antifade Mountant (Thermofisher), and images were taken using a Nikon C2 confocal microscope with 20× or 40× lenses as needed.

#### Calcium imaging

NGN2-Glut were replated on rat astrocyte coated coverslips on D5 DIV (days in vitro). On D35 DIV, neurons were labeled with 5 µM Fluo-4-AM at 37°C for 20 min. Coverslips were then transferred to a CO_2_ saturated 1× ACSF perfusion system for imaging. Excessive dyes were subsequently washed out by initial perfusion of 10-15 min. The time-lapse images were acquired at 1.8 Hz for 10 min on a Nikon A1R multiphoton microscope with a resonant scanner at 820 nm. For quantification, peak detection and amplitude measurement, regional intensity values were tabulated as one-dimensional numerical vectors indexed by time, and a Savitzky-Golay filter (locally estimated scatterplot smoothing) was applied for data smoothing. Baseline intensity level (for both peak detection and dF/F0 calculation) was performed using the R baseline package ^65^. Image analyses and quantification were performed in ImageJ and R. (See Supplementary Materials and Methods, and Supplementary Codes for all the parameters and programming details.)

#### Sholl and Puncta Analyses

For Sholl analysis, GFP-transfected neurons were imaged on a Nikon C2+ confocal microscope with a 10× objective lens and NIS-Elements software. Following the acquisition, dendrites were traced and images binarized in ImageJ. Sholl analysis was performed using the ‘Sholl analysis’ plugin for ImageJ. The center of the soma was manually defined, and concentric circles spaced 10 μm apart were used to quantify dendritic complexity in all cells.

For puncta and protrusion density analysis, GFP-transfected neurons were imaged using the same microscope with a 63× oil immersion objective lens. Dendritic branches were imaged on multiple z-planes to capture full dendritic branches, and maximum intensity projections of z-stacks created in ImageJ were used for downstream puncta analysis. Regions of interest (ROI) on the dendrite (∼100 μm) were selected using the GFP-filled cell as an outline. To identify puncta, a set threshold method was used for each experiment and applied identically to all conditions. The area and intensity of each punctum above the set threshold were recorded, and mean values were calculated for each cell. For puncta density analysis, the number of puncta was recorded and divided by the total area of the ROI calculated in ImageJ. For protrusion density analysis, a region of the dendrite (∼100 μm) was selected, and all protrusions <15 μm were manually counted and recorded.

For Sholl analysis, a two-way repeated measures ANOVA was performed to detect an effect of genotype-distance interaction. A *post hoc* correction was then applied to the analysis to correct for multiple testing and determine the significance of each data point at each distance interval. For synaptic puncta and protrusion analyses, data sets were tested for normality using D’Agostino and Pearson omnibus normality test. A one-way ANOVA (for parametric data) or a Kruskal Wallis test (for non-parametric data) with *post hoc* correction was used to determine significance between groups. All data were analyzed in GraphPad Prism version 9. Results were considered as significant if *p* < 0.05. All data are reported as mean ± SEM. None of the data was removed as outliers.

#### MEA Analysis

NGN2-Glut were prepared using NGN2-hygro virus as described in section “Generation of NGN2-Glut neurons”. Neurons were replated on Day 28 with rat astrocytes onto 0.1% PEI coated 24-well MEA plate (Axion BioSystems# M384-tMEA-24W). Media was refreshed every three days using Neurobasal Medium supplemented with 1× Glutamax/B27, 10 ng/ml BDNF/GDNF/NT-3, and 1% FBS. MEA data was recorded 24 hrs post media refreshment at 37°C with 5% CO_2_. On the day of recording, the plate was loaded into Axion Maestro MEA reader (Axion Biosystems) and allowed to rest for 3 min, then recorded for 10 min. Data files were batch-processed using Neural Metrics Tool (Axion Biosystems) with four biological replicates for each condition.

#### Statistical analysis

Unless otherwise specified, Student’s *t*-test (between two groups) or the Kruskal Wallis test with Dunn’s multiple comparisons and *p*-value adjustment (more than two groups) was used to determining significance between groups. Samples were assumed to be unpaired and have non-parametric distribution unless otherwise specified. Data were analyzed using R 4.1.1 and GraphPad Prism 9. Results were considered as significant if *p* < 0.05 (*: *P* < 0.05; **: *P* < 0.01; ***: *P* < 0.001; ****: *P* < 0.0001). All data are reported as mean ± SEM.

## Acknowledgements

We thank Rutgers University Cell and DNA Repository (RUCDR; under support of 2U24MH068457) for producing hiPSC lines. We thank Molecular Genetics of Schizophrenia (MGS) investigators for collecting samples that were used for hiPSC production. MGS includes P. V. Gejman, A. R. Sanders, J. Duan (NorthShore University HealthSystem, and University of Chicago, IL, USA), D. F. Levinson (Stanford University, CA, USA), J. Shi (National Cancer Institute, MD, USA), N. G. Buccola (Louisiana State University Health Sciences Center, LA, USA), B. J. Mowry (Queensland Centre for Mental Health Research, Brisbane and Queensland Brain Institute, The University of Queensland, Australia), R. Freedman, A. Olincy (University of Colorado Denver, CO, USA), F. Amin (Atlanta Veterans Affairs Medical Center and Emory University, GA, USA), D. W. Black (University of Iowa Carver College of Medicine, IA, USA), J. M. Silverman (Mount Sinai School of Medicine, NY, USA), W. F. Byerley (University of California at San Francisco, CA, USA), C. R. Cloninger, D. M. Svrakic (Washington University, MO, USA). We thank the study participants of MGS. MGS was mainly supported by R01MH059571, R01MH081800, and U01MH079469 (to P.V.G.); and other NIH grants for other MGS sites (R01MH067257 to N.G.B., R01MH059588 to B.J.M., R01MH059565 to R.F., R01MH059587 to F.A., R01MH060870 to W.F.B., R01MH059566 to D.W.B., R01MH059586 to J.M.S., R01MH061675 to D.F.L., R01MH060879 to C.R.C., U01MH046276 to C.R.C., and U01MH079470 to D.F.L). We thank Dr. Gejman for helping with the data interpretation. We also thank the PGC-SZ group for providing early access of wave-3 (PGC-SZ3) summary statistics.

## Author Contributions

S.Z. analyzed the ATAC-seq, bulk, and scRNA-seq data, performed most genetic/genomic analyses and wrote the manuscript. S.Z., H.Z. performed the experiments, analyzed data, and wrote the manuscript. M.P.F. performed immunostaining, imaging and analysis of dendritic and synaptic assays, gene set enrichment analysis, and wrote the manuscript. A.K. helped with neuron differentiation. V.A.B. and L.E.D. performed imaging and analysis of dendritic and synaptic assays. D.S. assisted with calcium imaging analysis. N.H.P. assisted with immunostaining and analysis. A.R.S. helped with clinical phenotypes, data interpretation, and manuscript writing. Z.P.P. guided the neuron differentiation. Y.F. helped with the CROP-seq data analysis. X.H. supervised the data analyses of Y.Z., and wrote the manuscript. P.P. supervised the neuronal phenotype analyses and wrote the manuscript. J.D. conceived the study, supervised the experiments and analyses, and wrote the manuscript.

## Data and Code Availability

The ATAC-seq and RNA-seq data is accessible under GSE188491. scRNA CROP-seq data is accessible under SRA PRJNA778419. All codes used in analysis is accessible at github.com/endeneon/VPS45_repo.

## Supplementary Materials and Methods

### hiPSC lines

The 20 hiPSC lines used for generating NGN2-Glut neurons for ATAC-seq were the same lines as used in our previously study ^13^. Briefly, hiPSCs were derived from cryopreserved lymphocytes (CPLs) using the genome-integration-free Sendai virus method (Cytotune Sendai Virus 2.0; Invitrogen). The purity of hiPSC culture was confirmed by immunofluorescence (IF) staining of pluripotency markers (OCT4, SSEA4, NANOG, and TRA-1-60). The pluripotency of the hiPSCs was further confirmed by Pluritest with RNA-seq data, as described in our previous study ^13^. Lines CD0000011 and CD0000012, which are heterozygous (A/G) at SNP site rs2027349, were used for CRISPR/Cas9 editing into all three genotypes, 2-3 clones per line ^13^. The NorthShore University HealthSystem Institutional Review Board (IRB) approved the study.

### hiPSC culture

The hiPSCs were maintained in mTeSR Plus media (STEMCELL) on matrigel (Corning) coated 6-well plate with media refreshment every other day. Cells were passaged at 1:10-1:20 every 4-6 days using ReLeSR (STEMCELL) following the vendor’s protocol.

### Generation of NGN2-Glut neurons

NGN2-Glut neurons were prepared as described ^20^ with modifications. Briefly, on Day 0, 5 × 10^5^ hiPSCs were dissociated using Accutase (STEMCELL) and replated in 1.2 ml mTeSR Plus media containing 5 µM ROCK inhibitor, rtTA virus and Ngn2-puro or Ngn2-Hygro virus on 6-well plates. On Day 1, media was refreshed with mTeSRPlus containing 2 µg/ml doxycycline. From days 2 to 4, media was refreshed daily with Neurobasal Medium supplemented with 1× Glutamax/B27, 2 µg/ml doxycycline, and 2 µg/ml puromycin or 200 µg/ml hygromycin. On day 5, cells were dissociated with Accutase and resuspended in Neurobasal Medium supplemented with 1× Glutamax/B27, 2 µg/ml doxycycline, and 10 ng/ml BDNF/GDNF/NT-3. Cells were replated at 2 × 10^5^/cm^2^ on 0.1% PEI/Matrigel coated 12-well plates. For morphological immunostaining, 5 × 10^4^ iNs were replated onto 12 mm coverslips (with 5 × 10^4^ rat astrocytes pre-cultured) in Neurobasal Medium supplemented with 1× Glutamax/B27, 2 µg/ml doxycycline, 10 ng/ml BDNF/GDNF/NT-3, and 1% FBS. From Day 6, media was refreshed every three days with a half volume change. Doxycycline was withdrawn from Day 14 onwards. ATAC-seq samples were harvested at Day 20; RNA-seq/qPCR samples were harvested at Day 30. For cells cultured on coverslips, GFP transfection was performed at Day 30 using Lipofectamine 3000 (Thermofisher) at 1:2 DNA:reagent ratio with 1 µg plasmid per coverslip. On Day 32, coverslips were fixed and stained for morphological analysis.

### Immunocytochemistry

Characterization of hiPSCs and neurons were performed as described ^13^ with minor modifications. Briefly, fresh cells were fixed in 4% PFA for 15 min at room temperature. After three PBS washes, cells were permeabilized by 1% Triton X-100 in PBS for 15 min at room temperature. Subsequently, samples were blocked with 3% BSA in 0.1% PBST (0.1% Triton X-100 in PBS) for one hour at room temperature. After blocking, samples were incubated with primary antibodies diluted in blocking buffer at 4°C overnight. After three PBS washes, samples were incubated with secondary antibodies diluted in blocking buffer for one hour at room temperature. After another three PBS washes, samples were incubated with 1 µg/ml DAPI for 10 min at room temperature. Samples were then mounted on glass slides using ProLong™ Diamond Antifade Mountant (Thermofisher) overnight before taking images. Primary antibodies used were anti-MAP2 (Synaptic System, 1:700), anti-vGlut1 (Synaptic Systems, 1:100), anti-GFP (chicken, Abcam ab13970, 1:10,000), anti-Synapsin I (rabbit, Cell Signaling #5297, 1:200), and anti-PSD-95 (mouse, NeuroMab clone K28/43 1:1000). Secondary antibodies used were Alexa 488 donkey anti-mouse (1:1000), Alexa 594 donkey anti-mouse (1:1000), Alexa 488 donkey anti-rabbit (1:1000), and Alexa 594 donkey anti-rabbit (1:1000). Images were taken using a Nikon C2 confocal microscope with 20× or 40× lenses as needed.

### ATAC-seq sample preparation

ATAC-seq samples were prepared as previously described ^13, 60^. Briefly, 75,000 viable cells were lysed in ATAC-Resuspension Buffer (RSB) containing 0.1% NP-40, 0.1% Tween-20, and 0.01% Digitonin for 3 min on ice. Nuclei were washed and resuspended in transposition mixture. Reactions were incubated at 37°C for 30 min on a thermomixer at 1,000 rpm. Transposed DNA was purified by DNA Clean and Concentrator-5 Kit (Zymo). Eluted DNA was shipped to the University of Minnesota Genomic Center for library preparation and ATAC-seq.

### ATAC-seq read aligning

All raw sequence reads generated by Illumina HiSeq 2000 had been demultiplexed at the University of Minnesota Genomics Center and provided as 2×75 bp paired-end fastq files (targeting 60 M reads per sample). Adapter remnants, low-quality reads, and low QSEQ short sequences near either end of reads were processed by Trimmomatic `(ILLUMINACLIP:NexteraPE-PE.fa:2:30:7, SLIDINGWINDOW:3:18, MINLENGTH:26)`. The processed sequences were separated into paired-end and single-end fastq files per sample and only paired-end reads were retained for subsequent mapping. The fastq files were individually mapped against the human genome reference file including decoy sequences (GRCh38p7.13/hg38, 1000 Genome Project) using bowtie2 `(-x 2000, -mm --qc-filter –met 1 –sensitive –no-mixed -t)` and subsequently merged and sorted as BAM-formatted files using samtools, with only uniquely mapped reads (MAPQ > 30) retained. Picard tools `MarkDuplicate` was then used to remove all PCR and optical duplicated reads from the BAM file.

To further eliminate allelic bias towards reference alleles during the aligning step, we performed WASP calibration on the generated raw BAM files ^61^. Briefly, we first called the VCF file profiles on all SNP sites that were not reference alleles of all 20 samples individually using GATK `HaplotypeCaller`, and subsequently collapsed the individual VCF files into one summary VCF file containing all non-reference sites of all 20 individuals. Subsequently, this SNP list was used as the basis of WASP calibration and re-alignment, and a new WASP-calibrated new BAM file set was collected as the final output for the following peak calling and ASoC SNP call ^61^.

The insert size distribution histograms of each sample were individually generated using Picard `CollectInsertSizeMetrics`. All analyzed ATAC-seq samples passed standard QC based on the characteristic nucleosomal periodicity of the insert fragment size distribution and high signal-to-noise ratio around transcription start sites (TSS).

### ATAC-seq peak calling and ASoC SNP calling

To increase sample size and sensitivity for peak detection, the BAM files of the processed reads of each sample were first sub-sampled to 30 M pair-end reads per sample (the smallest sample size) and the 20 sub-sampled BAM files were merged as the source file for peak calling. MACS2^25^ was used to generate peak files (narrowPeak format) with recommended settings at `FDR = 0.05 (-f BAMPE, --nomodel, --call-summits --keep-dup-all -B)`. Peaks that fell within the ENCODE blacklisted regions were removed. Also, we removed peaks falling within chromosomes X and Y, and the mitochondrial genome regions.

GATK (version 4.1.8.1) was used for ASoC SNP calling, as recommended by the GATK Best Practices (software.broadinstitute.org/gatk/best-practices/) ^62^. As noted above, WASP-calibrated BAM files (without sub-sampling) generated from ATAC-seq pipeline were used as input. Variants were called using the discovery mode of `HaplotypeCaller` with human GRCh38 (hg38) genome and the corresponding dbSNP version 153, and only reads with MAPQ score ≥30 were used `(-stand_call_conf 30)`. Subsequently, recalibration of SNPs and Indels were performed in tandem using the `VariantRecalibrator` function `(-an DP -an QD -an FS -an SOR -an MQ - an ReadPosRankSum -mode SNP -tranche 100.0 -tranche 99.9 -tranche 99.5 -tranche 90.0 -mG 4)` and applied using `ApplyRecalibration. Databases` used for `VariantRecalibrator` including the hg38 versions of HapMap v3.3 (priority = 15), 1000G_omni v2.5 (priority = 12), Broad Institute 1000G high confidence SNP list phase 1 (priority = 10), Mills 1000G golden standard INDEL list (priority = 12), and dbSNP v153 (priority = 2). Each sample within the cell type was processed individually, and heterozygous SNP sites with tranche level >99.5% were extracted. To reduce bias introduced by any acquired (or “*de novo*”) mutations during cell growth, only SNPs with corresponding rs# records found in dbSNP v153 were retained. All heterozygous sites that passed the filter above (20 samples) were merged by CombineVariants to produce the master VCF file of the cell type.

To maximize the power to detect ASoC, we pooled SNPs for all the called heterozygous SNPs locations from the 20 samples as justified by the high concordance of allele-specific effects across individuals ^13^. Finally, the VCF files were filtered and only biallelic SNP sites (GT: 0/1) with minimum read depth count (DP) ≥ 20 and minimum reference or alternative allele count ≥ 2 were retained. The binomial *p*-values (non-hyperbolic) were calculated using the `binom.test(x, n, P = 0.5, alternative = “two.sided”, conf.level = 0.95`) from the R package, and Benjamini & Hochberg correction was applied to all qualified SNPs as the correcting factor of R function `p.adjust(x, method = "fdr")`. We set the threshold of ASoC SNP at FDR value = 0.05.

The read pileup statistics proximal to SNP sites were generated using `samtools mpileup` function, and differential of allele-specific reads was performed using the SNPsplit Perl package (www.bioinformatics.babraham.ac.uk/projects/SNPsplit/). The final readouts from both read pileup and SNP-specific reads were visualized using the R package Gviz. In addition, when comparing the changes of chromatin accessibility caused by genotypes across samples, read counts were scaled and normalized using the deepTools package bamCoverage function and re-scaled to reads per genomic content (RPGC) as the base unit ^63^. We confirmed no obvious mapping bias to reference alleles by visualizing the volcano plots that graph the allelic read-depth ratios against -log_2_*p*-values in scatter plots (Extended Data Fig. 2a).

### Principal component analysis (PCA) for ATAC-seq data

To compare the cellular property between our NGN2-Glut neurons and different hiPSC-derived neuronal cell lines generated in our lab, as well as with previously assayed fetal brain cells, we used chromatin accessibility data from our previously published studies ^13^ as well as DNase I hypersensitive assay from the fetal brain (NIH Roadmap) ^64^. Subsequently, we applied our above described cross-cell-type non-overlapping OCR peak interval (n = 666,614) (GTF file, genome lift-over was performed when necessary) to the 20 lines of NGN2-Glut data using featureCounts^66^ to get per-interval read count. PCA was applied to the assembled datasets, integrating our NGN2-Glut into the multitype dataset (Extended Data Fig. 1g).

### Bulk RNA-seq and differential expression (DE) analysis

Total RNA was isolated using RNeasy plus kit (QIAGEN) following the vendor’s instructions and quantified by a NanoDrop-8000 spectrometer. The isolated RNA was subsequently sent to the sequencing provider (GENEWIZ) for further processing and sequencing analysis. RNA-seq files were provided in the format of 2×150 bp paired-end fastq files. Briefly, libraries were prepared using the NEB Nextera kit with customized adapters. Since the sequencing facility had performed pre-cleaning on raw reads, no Trimmomatic adapter-trimming was performed. 20∼30 M reads were recovered from each sample. Raw files were subsequently mapped to human hg38 genome (GRCh38p7.13) using STAR v2.7.0 ^57^ with the following parameters: (--outSAMtype BAM SortedByCoordinate --quantMode GeneCounts -- outSAMattrIHstart 0 --outSAMstrandField intronMotif --outSAMmultNmax 1 -- outFilterIntronMotifs RemoveNoncanonical --outBAMcompression 10 -- outBAMsortingThreadN 20 --outBAMsortingBinsN 20 --outFilterMultimapNmax 1 -- outFilterMismatchNmax 1 --outSJfilterReads Unique --limitBAMsortRAM 10000000000 --alignSoftClipAtReferenceEnds No -- quantTranscriptomeBAMcompression 10 10).

PCR and optical duplicates were further removed using Picard MarkDuplicates, and unique mappers (MAPQ = 255) were retained for downstream analysis. GENCODE v35-based gene annotations, transcript length, and GC percentage were used ^67^. For gene-based quantifications, the number of fragments at the meta-feature (gene) level was directly collected from the output of STAR, and only genes expressed in at least one cell type with counts per million (CPM) >1 in at least half of the sample within the group were retained for subsequent analysis. CPM numbers were used to plot the correlation heatmap and PCA analysis. For transcript-based quantifications (such as in the case of quantifying the expression level of different *VPS45* isoforms), the pseudo- alignment probabilistic model of Kallisto was applied ^58^. The PCA map was generated using R packages, gplots and ggplot2, respectively. DE gene analysis was performed using EdgeR ^59^ by fitting the dataset into a gene-wise negative binomial generalized linear model with quasi-likelihood as glmQLFit() and evaluated by `glmQLFTest()`. Cell line and batch numbers were introduced when establishing the design matrix and served as blocking factors, whenever applicable.

CPM values generated by EdgeR was used as the basis for making the PCA plot to visualize the clustering effect of different gene knockdown groups. Essentially, only the top 15,000 DE genes were used for calculating the PCs using R `prcomp()` function and the plot was subsequently generated using the `fviz_pca()` function from R package `factoextra` (Extended Data Fig. 9b).

### Total RNA extraction and qPCR analysis

For bulk RNA-seq and qPCR, RNA was isolated using RNeasy plus kit (QIAGEN) following the vendor’s instructions and quantified using NanoDrop-8000 spectrometer. For qPCR, RT was performed using High-Capacity cDNA Reverse Transcription Kit (Thermofisher). RT conditions were as follows: 25°C for 10 min, 37°C for 120 min, 85°C for 5 min. qPCR was performed on a Roche 480 II Lightcycler, using gene-specific FAM-labelled TaqMan probes, *GAPDH*, *ABL1* and *CASC3* as internal controls. qPCR conditions were as follows: 95°C for 10 min, 45 cycles of (95°C for 20 s, 60°C for 1 min). Biological replicates (3 to 6) were included for all conditions and three technical replicates were used during qPCR.

### Gene expression correlation between DE genes from SNP editing and for various disorders

The correlation of expression profiles of DE genes between *VPS45* CRISPR/Cas9-edited lines (A/A vs.G/G) and postmortem brain tissues of various psychiatric disorders (MDD, BP, ASD, AAD, SZ) were compared by integrating GTEx/PsychENCODE and several other data sources ^15, 68, 69^. For each DE gene set, genes with FDR < 0.05 were used in the overlapped gene counts and correlation analysis. The R package `upsetR` was used to make the Upset plot. For correlation analysis, Spearman’s ρ was calculated using R function `cor.test(method = “spearman”, alternative = “t”)`.

### Lentivirus preparation

Lentivirus particles were packaged in fresh HEK 293T culture (passaged ≤ 15 times) as previously described ^13^. Briefly, 2 × 10^6^ cells were replated in T25 flasks one day before transfection. At the day of transfection, media was refreshed without antibiotics. pLenti-dCas9-KRAB-BSD (Addgene # 89567) was co-transfected with pMD2.G (Addgene #12259) and psPAX2 (Addgene #12260) at a 4:2:3 molar ratio; other viral vectors including FUW-M2rtTA (Addgene # 20342), pTet-O-Ngn2-puro (Addgene # 52047), CROPseq-Guide-Puro gRNA (Addgene # 86708), and pTet-O-Ngn2-hygro were co-transfected with pMDLg/pRRE (Addgene #12251), pMD2.G (Addgene #12259), and pRSV-Rev (Addgene #12253) at 1:1:1:1 molar ratio. Transfection was carried out using FuGENE HD (Promega) following the vendor’s instructions. 24 hrs post transfection, media was refreshed with Neurobasal Medium supplemented with 1× Glutamax/B27. 48 hrs post transfection, the supernatant was collected and centrifuged at 500 × g for 5 min to remove debris. The virus-containing media was aliquoted into 1.5 ml low protein binding tubes and stored at -80°C until use.

### CROP-seq in NGN2-Glut

hiPSC lines (Line CD000009 and CD000011) that stably express dCas9-KRAB-BSD were established as previously described ^13^. Briefly, on day 0, hiPSCs were replated as small clumps using ReLeSR (STEMCELL) at 10-20% confluence. On Day 1, refresh media with mTeSR Plus containing dCas9-KRAB-BSD virus. 24 hrs post transduction, media was refreshed with mTeSR Plus containing 5 µg/ml Blasticidin. Antibiotics selection was refreshed daily and maintained for 10 days.

NGN2-Glut were prepared from hiPSCs stably expressing dCas9-KRAB-BSD using NGN2-hygro virus as described in a previous section (Generation of NGN2-Glut neurons). From Days 2-4, cells were selected using 150 µg/ml hygromycin (Sigma). On Day 20, iNs were dissociated using Accutase and replated in Neurobasal Medium supplemented with 1× Glutamax/B27, 10 ng/ml BDNF/GDNF/NT-3, 5 µM ROCK inhibitor, and CROPseq-Guide-Puro virus (vector carrying gRNA targeting genes of interest, MOI=0.2) as described in the original CROP-seq article ^28^. 48 hrs post transduction, cells were selected using 2 µg/ml puromycin. 7 days after selection, cells were harvested using Accutase for single cell RNA-seq preparation using Chromium Next GEM Single

### Single-cell RNA-seq data processing and analysis

10× Genomics Cell Ranger v4 was used to process raw sequencing data. Briefly, sequencing data were aligned to the human GRCh38/hg38 genome with spike-in gRNA sequences as artificial chromosomes (20 bp gRNA sequence and 250 bp downstream plasmid backbone per each gRNA, together with one additional full sequence of CROP-Seq-Guide-Puro construct also included as a decoy to capture vector-borne sequences). The alignment was performed using STAR 2.7.0 included in Cell Ranger v4 and a customized spiked-in version of GENCODE v28 GTF file containing the aforementioned gRNA sequences and vector backbone as annotations. The filtered gene count matrix output from Cell Ranger was used in subsequent R analysis. In all, we achieved a total mapping rate of 88.7%, with 58.1% mapping rate to the human transcriptome.

The R package Seurat v 3.2 was used to extract information and assemble data tables from the output files to generate the digital gene expression matrix, which was constructed based on per-gene UMI count. Since the Cell Ranger package had already performed preliminary QC filtering and removed low-counting barcodes (*i.e.*, cells) from its filtered output data set (`filtered_bc_matrix`), we directly imported the output of these processed barcode matrices as raw data. Outputs from one capture (using a mix of CD000009 + CD000011) were imported, which resulted in a total of 10,247 cells used in the initial analysis. The Seurat matrix was subsequently re-normalized using `SCTransform()` and analyzed on the generated SCT object. Cells with > 20% mitochondria content were removed from analysis after SCTransform. A cell assigned with a unique RNA was defined as the UMI count of its dominant gRNA in the cell being at least three times more than the sum of UMI counts from all other gRNAs. Only MAP2+ cells that express glutamatergic neuron marker genes (SLC17A6 or SLC17A7) falling within clusters 2, 3, 4, 5, 6, and 7 were used (Fig. 2b). Eventually, 4,057 cells survived the filter and were designated as cells with uniquely assigned gRNA for subsequent study. Dimension reduction, UMAP, clustering, and pseudo-bulk DE analysis were performed using package Seurat and EdgeR. Graphs were made in Seurat and ggplot2.

### MAGMA analysis

We performed MAGMA analysis using MAGMA version 1.08b ^70^ to evaluate the enrichment for the GWAS risk of several psychiatric disorders (SZ, Neuroticism, ASD, BD, AAD, MDD) ^1, 4, 5, 40–42^, as well as the GWAS set of Crohn’s disease that served as a control set ^43^. Specifically, we first started by compiling the MAGMA-required gene annotation data files using an GRCh37/hg19 vcf file. With the gene-SNP annotation file, we then performed gene-level analysis on SNP *p*-values using the reference SNP data of 1,000 Genomes European panel (g1000_eur, `--bfile`) and the pre-computed SNP *p*-values from each disorder’s GWAS data set. The sample size (`ncol=`) was also either directly taken from the column of sample sizes per SNP column of the datasets or extracted from the affiliated README data. Subsequently, the result files (`--gene-results`) from the gene-level analysis were read in for competitive gene-set analysis (`--set-annot`), where we used default setting (`‘correct=all’`) to control for gene sizes in the number of SNPs and the gene density (a measure of within-gene LD). The gene-set analysis produced the output files (`.gsa.out`) with competitive gene-set analysis results that contained the effect size (BETA) and the statistical significance of the enrichment of each gene set (All DE genes/upregulated/downregulated) for each disorder’s GWAS data set. The heatmap graph was plotted using R package `gplots`.

### Gene set over-representation analysis (ORA) with WebGestalt

We performed ORA using WebGestalt ^34^. Briefly, lists of DE genes from RNA-seq analysis were input into the web-based interface of WebGestalt to output the enriched (or over-represented) GO-terms. All the expressed genes in NGN2-Glut were used as a reference gene list for the enrichment analysis. The GO-terms included three different categories: biological processes, cellular components, and molecular function. The fold of enrichment and the -log_10_ *p*-value for the enriched GO-terms were plotted as volcano plots, which were outputted directly from the browser and then polished and re-arranged for clarity purposes in Adobe Illustrator.

### SynGO GE and ontology analysis

SynGo GE and ontology analysis were performed as documented ^71^ using the “location” domain and colorized by either “enrichment analysis” or “gene count”, as indicated on the generated Sunburst plots. Briefly, the list of DE genes was used as the SynGO input list, and all the expressed genes in NGN2-Glut neurons were used as the background gene list. The exact gene list per term was further extracted using the select term function and exported for annotation.

### Gene set enrichment analysis

Lists of DE genes were divided into three subsets: all dysregulated genes, upregulated and downregulated (all at FDR < 0.05). DE genes were overlapped with gene sets relevant to SZ, including genes implicated by GWAS and the post synaptic density proteome ^27, 72, 73^. The GWAS list included two subsets: “Single genes” subset included all GWS loci implicating single genes, and a second, larger subset combining all genes within single-gene or multigenic GWAS loci. A hypergeometric test was used to calculate the extent of over- or under-enrichment of disorder-relevant genes compared to a chance finding. All genes expressed genes in NGN2-Glut neurons were used as a background set.

### Gene knockdown (KD) in NGN2-Glut

Short hairpin RNA (shRNA) sequences were designed using Thermofisher online designer (rnaidesigner.thermofisher.com/rnaiexpress/insert.do) following antisense-loop-sense pattern. Refer to Table S15 for shRNA sequences. Single-strand shRNA oligos were cloned into BsmBI digested CROPseq-Guide-Puro (Addgene# 86708) vector using NEBuilder® HiFi DNA Assembly Master Mix (New England Biolabs) following vendor’s protocols. Oligo sequence targeting GFP was served as control. Lentivirus particles were prepared as described in previous sections (Lentivirus preparation). Stable knockdown hiPSC lines were established as follows. On Day 0, hiPSCs were replated as small clumps using ReLeSR (STEMCELL) at 10-20% confluency. On Day 1, media was refreshed with mTeSRPlus containing CROPseq-KDgRNA-Puro virus. 48 hrs post-transduction. Media was refreshed with mTeSRPlus containing 0.8 µg/ml puromycin. Antibiotics selection was refreshed daily and maintained for 10 days. NGN2-Glut were generated from Ngn2-hygro lentivirus as described in a previous section (Generation of NGN2-Glut). Total RNA was isolated from DIV30 cultures using RNeasy Plus kit (QIAGEN) for qPCR quantification and RNA-seq.

### Comparison of shRNA KD results between the two cell lines

In order to compare the concordance of shRNA KD behavior between the two independently generated, SNP-edited A/A (rs2027349) cell lines (A11, H12), we performed RNA-seq using shRNA targeting *EGFP, VPS45, AC244033.2*, and *C1orf54* on the two cell lines. For each line, three biological replicates were used, and the DE gene list generated within each line was paired with the corresponding DE gene list from the other line to form a union set. The union set of genes of each KD condition was used as the index for the corresponding log_2_FC values in either line for regression analysis. Pearson’s R was used to show the correlation between the log_2_FC values from different lines.

### Linear regression analysis to correlate the log_2_ fold changes (FC) of DE genes from SNP editing and their log_2_FC in different shRNA KD conditions

To evaluate which individual gene KD and/or their interaction terms could better explain the transcriptional effects of rs2027349 (*VPS45*) editing in NGN2-Glut, we employed a general linear regression model to correlate the log_2_FC of DE genes in G/G neurons (vs A/A) and their log_2_FC under different shRNA KD conditions in A/A neurons. Because A/A neurons showed higher expression of the three local genes of interest (*VPS45, AC244033.2*,, *and C1orf54*) at the rs2027349 locus as a result of rs2027349 editing, if a gene or gene × gene interaction could explain or mediate the transcriptional effect of the SNP editing, we would expect a significant correlation between log_2_FC of DE genes in G/G neurons (vs A/A) and their log_2_FC in a gene KD, or a combination of KD of two genes in A/A neurons. Because of the noisy nature of the gene KD experiment that showed an excessive number of DE genes, we only analyzed the list of 1,267 significant DE genes in G/G neurons (vs A/A). We included individual gene KD and their interaction terms in the following model:

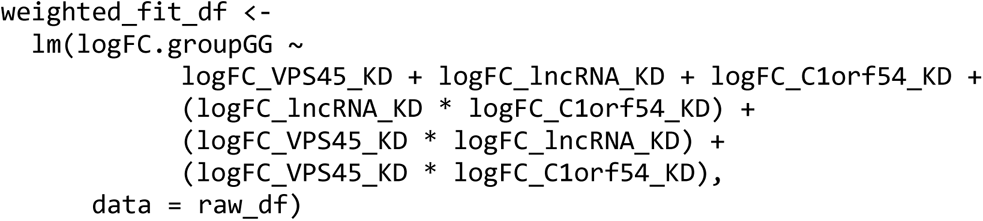

in which raw_df is the data frame that contained all DE expression data. The derived beta coefficient and *p*-value were used to infer the association of SNP editing effects and individual gene KD and their interaction terms.

### Sholl analysis

GFP-transfected neurons were imaged on a Nikon C2+ confocal microscope with a 10× objective lens and NIS-Elements software. Neurons were only imaged if a visible axon and at least one defined primary dendrite could be identified. Following the acquisition, dendrites were traced and images binarized in ImageJ (National Institutes of Health, Bethesda, MD). For each image, the axon was identified and eliminated from quantification. Sholl analysis was performed using the ‘Sholl analysis’ plugin for ImageJ. The center of the soma was manually defined, and concentric circles spaced 10 μm apart were used to quantify dendritic complexity in all cells.

### Puncta and protrusion density analysis

GFP-transfected and immunostained neurons were imaged using a 63× oil immersion objective lens on a Nikon C2+ confocal microscope with NIS-Elements software. Each channel was acquired sequentially to prevent fluorescence bleed-through with 2× line averaging for each channel and 1.5× digital zoom. Image parameters were optimized for each batch (*i.e.*, all neurons from the same differentiation) to limit the over- or undersaturation of signal. All images within a batch were acquired using identical settings for each channel to allow comparison between groups. Dendritic branches were imaged on multiple z-planes to capture full dendritic branches and maximum intensity projections of z-stacks created in Image J were used for downstream puncta analysis. Regions of interest (ROI) on the dendrite (∼100 μm) were selected using the GFP-filled cell as an outline. To identify puncta, a set threshold method was used for each experiment and applied identically to all conditions. The area and intensity of each punctum above the set threshold were recorded, and mean values were calculated for each cell. For puncta density analysis, the number of puncta was recorded and divided by the total area of the ROI calculated in ImageJ. For protrusion density analysis, a region of the dendrite (∼100 μm) was selected, and all protrusion <15 μm were manually counted and recorded.

### MEA Analysis

NGN2-Glut were prepared as described in a previous section. On Day 28, neurons were dissociated using Accutase and replated at 1 × 10^5^ cells/well with 1.5 × 10^4^ cells/well rat astrocytes onto a 0.1% PEI coated 24-well MEA plate (Axion BioSystems# M384-tMEA-24W). Media was refreshed every three days using Neurobasal Medium supplemented with 1× Glutamax/B27, 10 ng/ml BDNF/GDNF/NT-3, and 1% FBS. MEA data was recorded 24 hrs post media refreshment at 37°C with 5% CO_2_. On the day of recording, the plate was loaded into Axion Maestro MEA reader (Axion Biosystems) and allowed to rest for 3 min, then recorded for 10 min. Data files were batch-processed using Neural Metrics Tool (Axion Biosystems) with four biological replicates for each condition. For analysis, bursts were identified using an inter-spike interval (ISI) threshold requiring a minimum number of 5 spikes with a maximum ISI of 100 msec. Network bursts were analyzed using the envelope mode with threshold factor 1.25, minimum inter-burst interval (IBI) 100 msec, minimum of 25% active electrodes, and 75% burst inclusion. The synchrony index was calculated using a cross-correlogram synchrony window of 20 msec.

### Calcium imaging

NGN2-Glut were replated on rat astrocyte coated coverslips on D5 DIV. On D35 DIV, neurons were labeled with 5 µM Fluo-4-AM at 37°C for 20 min. Coverslips were then transferred to CO_2_ saturated 1× ACSF (126 mM NaCl, 2.5mM KCl, 1 mM NaH_2_PO_4_, 26.2 mM NaHCO_3_, 2.5 mM CaCl_2_, 1.3 mM MgSO_4_, 11 mM D-Glucose) perfusion system for imaging. After 10-15 min perfusion to wash out extra dyes, time-lapse images were acquired at 1.8 Hz for 10 min using a Nikon A1R multiphoton microscope at 820 nm. Quantified regional intensity values were tabulated as one-dimensional numerical vectors indexed by time, and a Savitzky-Golay filter (locally estimated scatterplot smoothing) was applied for data smoothing. Baseline intensity level (for both peak detection and dF/F0 calculation) was performed using the R baseline package ^65^. The maximum peak intensities were subsequently captured by detecting the maximum values on the rolling margins of each peak window (see the Extended Data code), and frequencies (the inversion of firing rate) were calculated using their corresponding peak positions. Amplitude (dF/F0) was calculated using the standard method by dividing the calibrated peak value at any specific position over the baseline intensity value at the same position as determined by the baseline package, and for each cell, one dF/F0 was calculated by taking the means of the dF/F0 values from all peaks from the same cell. Synchrony index was calculated using the original spike train distance model ^74^ by spike distance and temporal distance normalized as mean resultant length for inter-sample measuring and visualization. Images analysis and quantification were performed in ImageJ and R.

### Statistical analysis

Unless otherwise specified, Student’s t-test (between two groups) or the Kruskal Wallis test with Dunn’s multiple comparisons and *p*-value adjustment (more than two groups) was used to determining significance between groups. Samples were assumed to be unpaired and have non-parametric distribution unless otherwise specified. Data were analyzed using R 4.1.1 and GraphPad Prism 9. Results were considered as significant if *p* < 0.05 (*: *P* < 0.05; **: *P* < 0.01; ***: *P* < 0.001; ****: *P* < 0.0001). All data are reported as mean ± SEM.

---

**Extended Data Fig. 1:**
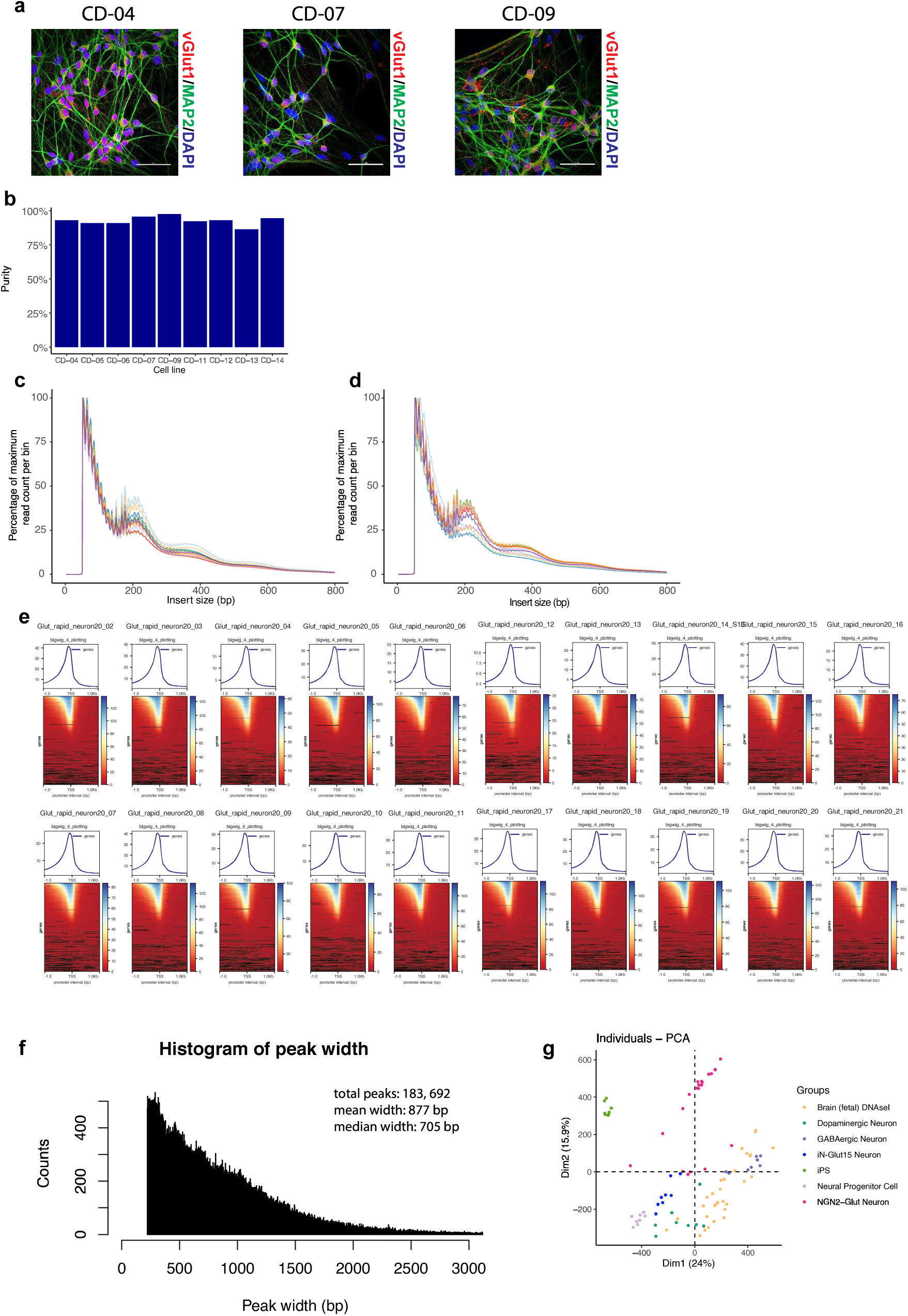
Morphological characterization and QC of hiPSC-derived NGN2-Glut neurons. (a) Representative immunofluorescence (IF) images of several NGN2-Glut lines stained with antibodies against glutamatergic cell-type-specific markers. Scale bar: 50 µm. (b) Cell line purity statistics, shown are the percentages of NGN2-Glut cells that are stained positive with type-specific markers (vGLUT1+/MAP2+). (c-d) Histogram of the insert sizes of all ATAC-seq samples used, showing the expected distribution pattern of transposase insertion at specific sizes (in bp; *x*-axis). 20 NGN2-Glut samples were plotted in two groups (10 each) for better visual resolution. (e) Chromatin accessibility in regions proximal to the TSS of core promoters (± 1 kb) in each sample, showing the library size-normalized overall ATAC-seq read intensity (top) and per-gene heatmap (bottom). The strong signal to noise ratio of the ATAC-seq peak intensity around the TSS site indicated a good quality of ATAC-seq data. Color bar indicates the density of reads at each position proximal to the TSS site. Black regions represent the absence of signal at the corresponding coordination. (f) The histogram of all ATAC-seq peak distribution. *x*-axis: peak width in bp. (g) PCA of OCRs from taken from NGN2-Glut, iPSC, the four iNs previous reported ^13^, and fetal brain DNAseI results, showing the relative distribution of NGN2-Glut neuron against other cell types.

**Extended Data Fig. 2:**
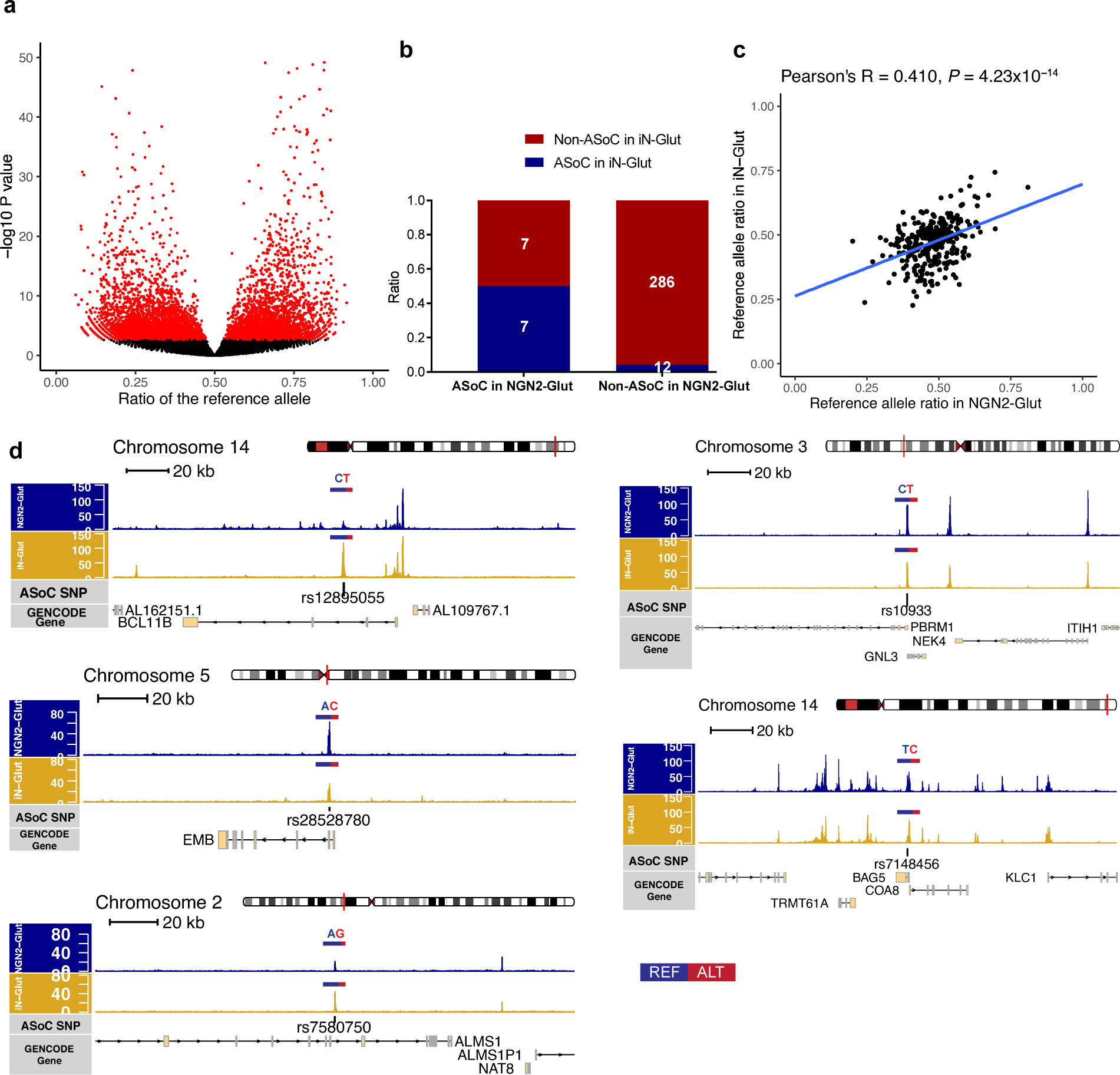
Allele-specificity in NGN2-Glut neurons. (a) Volcano plot showing the reference allelic ratio of all heterozygous SNPs (*x*-axis) and their corresponding *p*-values of ASoC testing (*y*-axis; in -log_10_ scale) from 20 NGN2-Glut samples. Red: all ASoC SNPs (FDR < 0.05). (b) Bar graph depicting the enrichment of overlapping ASoC SNPs associated with SZ in NGN2-Glut and iN-Glut data sets. Chi square test, *p* < 0.001. (c) Allelic ratio correlation of SZ credible SNPs between NGN2-Glut and iN-Glut neurons. (d) ATAC-seq read pileup and gene tracks proximal to five SZ-credible SNP sites in NGN2-Glut and iN-Glut neurons. the ratio of reference and alternate allele has been marked over each SNP as the length of color bars, showing high concordance between the two neuron types.

**Extended Data Fig. 3:**
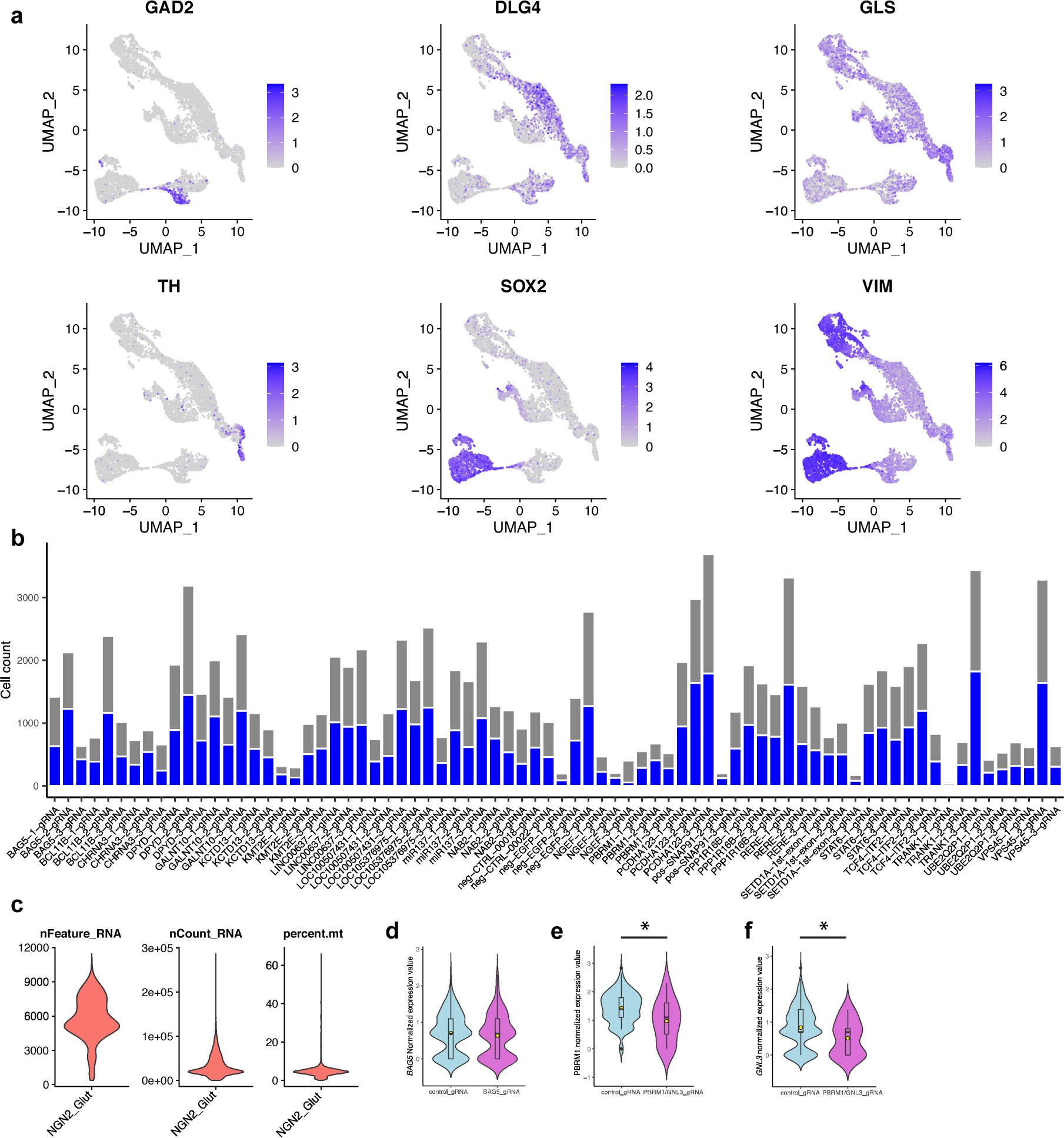
Quality control summary of CROP-seq experiment in NGN2-Glut neuron. (a) Expression feature plots showing additional cell type-specific markers from the output of scRNA-seq result. (b) Bar graph showing the number of cells (*y*-axis) that express a specific gRNA (*x*-axis) and their gRNA contents. Blue bar: cells were assigned with a unique gRNA and used for differential gene expression analysis. Five negative control gRNAs were used (neg_CTRL_0018, neg_CTRL_0022, neg_EGFP_1, neg_EGFP_2, and neg_EGFP_3). (c) General QC data output from Seurat after merging the two capture data sets and performed data normalization with SCTransform(). (d) Quantification on the knockdown efficiency of *BAG5* (rs7148456) CROP-seq gRNA on the expression level of *BAG5* gene. Student’s t-test (non-parametric). (e-f) Quantification on the knockdown efficiency of *PBRM1*/*GNL3* (rs10933) CROP-seq gRNA on the expression level of either *PBRM1* or *GNL3* gene. Student’s t-test (two-sided, non-parametric). For the control group, all five negative control gRNAs were used in quantification.

**Extended Data Fig. 4:**
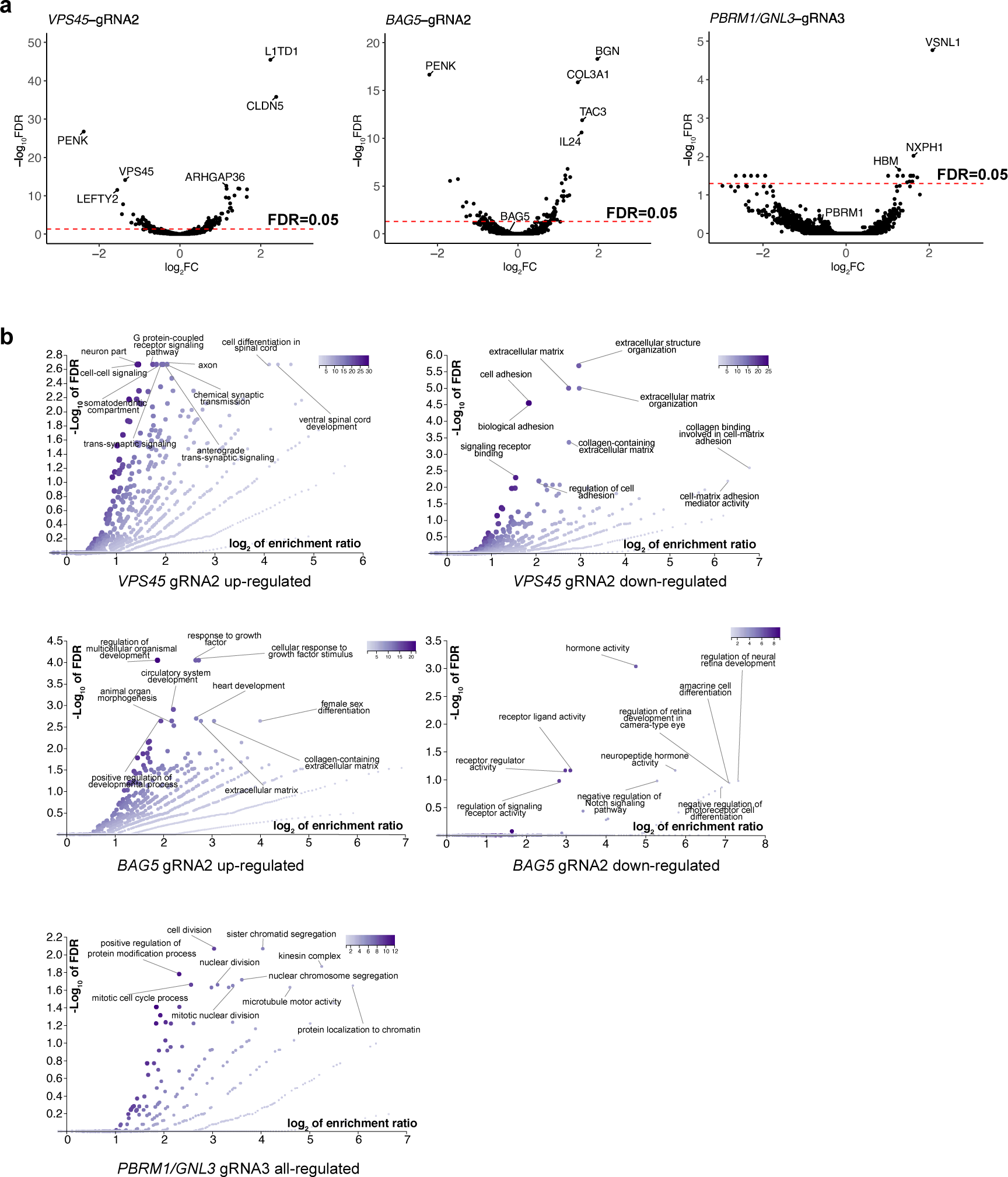
DE analysis of CROP-Seq result. (a) Volcano plots showing DE genes in different CROP-seq gRNA-containing cells, VPS45-gRNA2 (rs2027349), BAG5-gRNA2 (rs7148456), and PBRM1/GNL3-gRNA3 (rs10933). (b) GSEA for DE genes in CROP-seq at *VPS45*, *BAG5*, and *PBRML1*/*GNL3* loci. Plotted are fold enrichment and -log_10_ *p*-value for GO terms.

**Extended Data Fig. 5:**
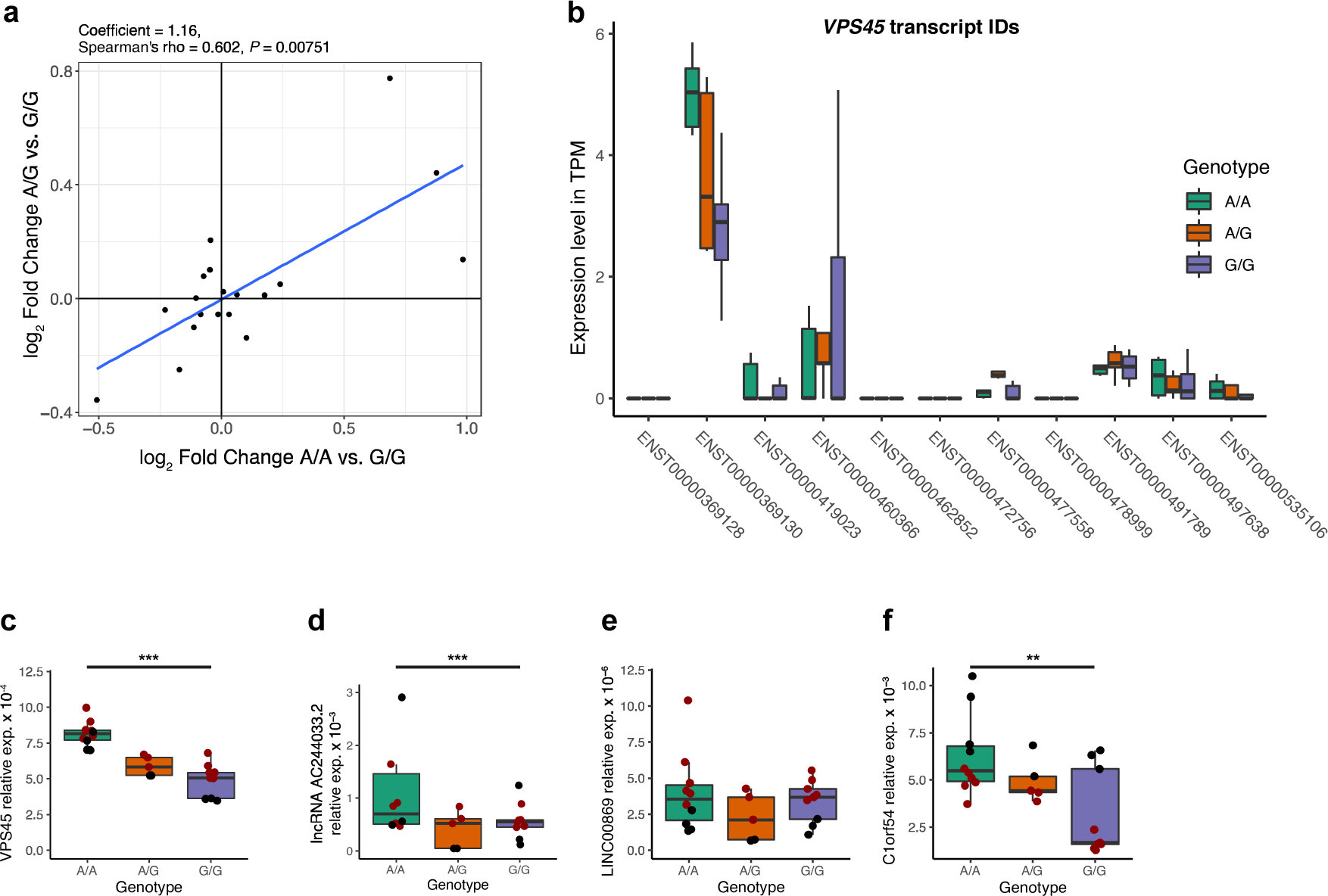
*Cis*-effects of SNP editing at the *VPS45* locus in NGN2-Glut of the rs2027349-edited isogenic lines. (a) *Cis*-effect with 500kb interval of the edited SNP site (rs2027349), demonstrating a high correlation on the DE gene expression patterns between AA/GG vs. AG/GG. (b) Comparison of the expression level of different *VPS45* transcripts in rs2027349 edited NGN2-Glut neurons from RNA-seq. Note the high expression level of main transcript ENST00000369130. (c-f). qPCR validation of VPS45-proximal DE genes from discovered from RNA-seq results. Points with different color indicate biological repeats from independent experiment groups. (c) VPS45. (d) AC244033.2. (e) LINC00869. (f) C1orf54. Kruskal-Wallis test (non-parametric).

**Extended Data Fig. 6:**
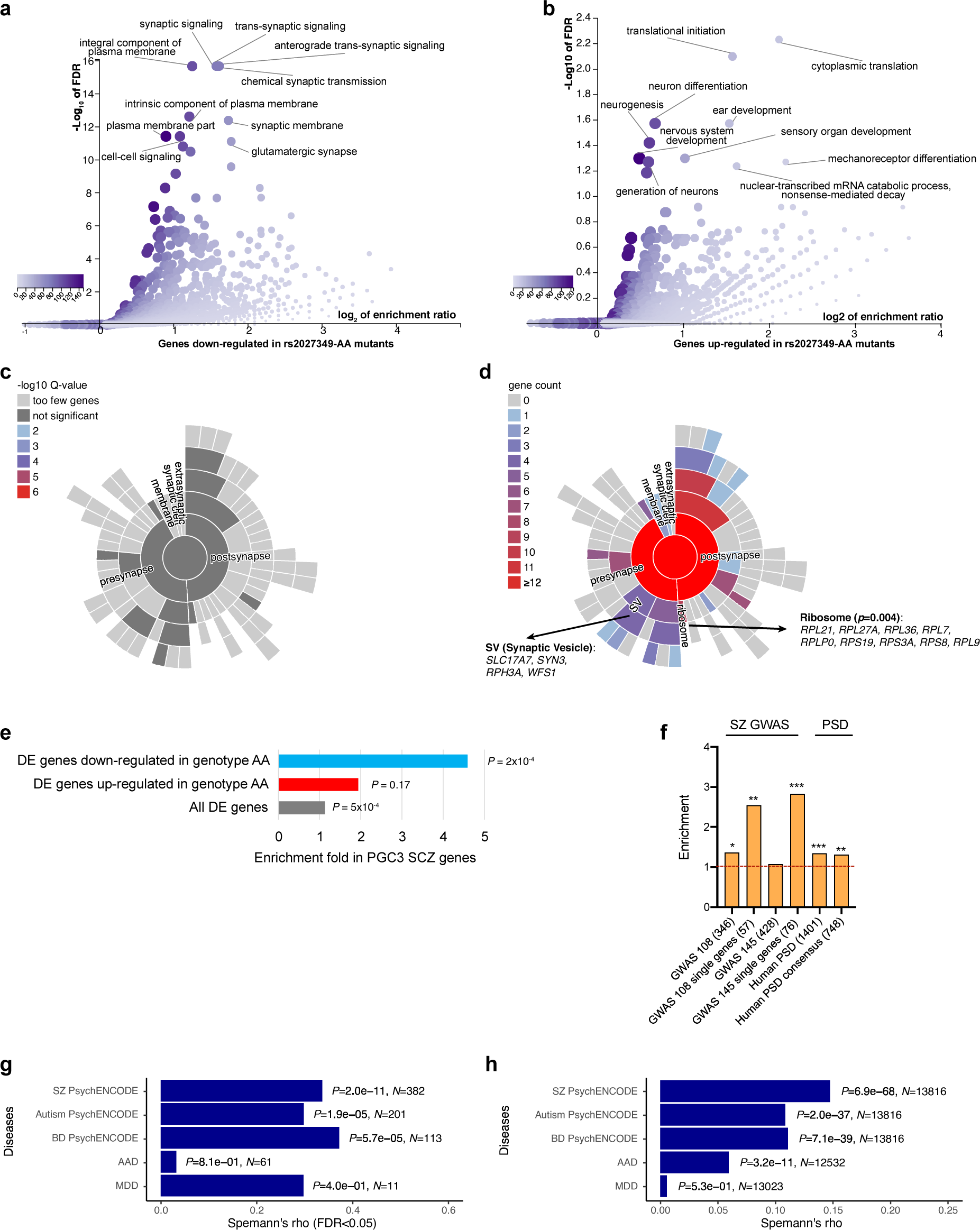
Transcriptomic gene pathway and disease relevance analyses in NGN2-Glut of the rs2027349-edited isogenic lines. (a-b) GO term analysis of DE genes in rs2027349-edited neurons. Genotype G/G was used as the expression baseline. (a) Genes downregulated in rs2027349-edited A/A neurons. (b) Genes upregulated in rs2027349 A/A neurons. (c) Sunburst plot showing the enrichment, albeit insignificant, of synaptic-related terms among the upregulated DE genes in rs2027349 A/A neurons. (d) Further dissection of (c) using the gene count as the coloring scheme, showing an excess number of synaptic vesicle and ribosome-related terms. (e) GSEA result of DE genes in rs2027349 A/A neurons for PGC3 SZ GWAS risk genes. (f) Enrichment of DE genes in rs2027349 A/A neurons for several SZ-related gene lists. Fisher’s Exact Test was used for the enrichment analyses in (e) and (f). (g) Correlation of the log_2_FC of DE genes (FDR < 0.05) from rs2027349 editing (A/A vs. G/G neurons) in NGN2-Glut neurons against the overlapping DE genes (FDR < 0.05) in postmortem brains of multiple neuropsychiatric disorders. (h) Similar to (g), except using all the expressed genes. Spearman’s ρ test, two-sided.

**Extended Data Fig. 7:**
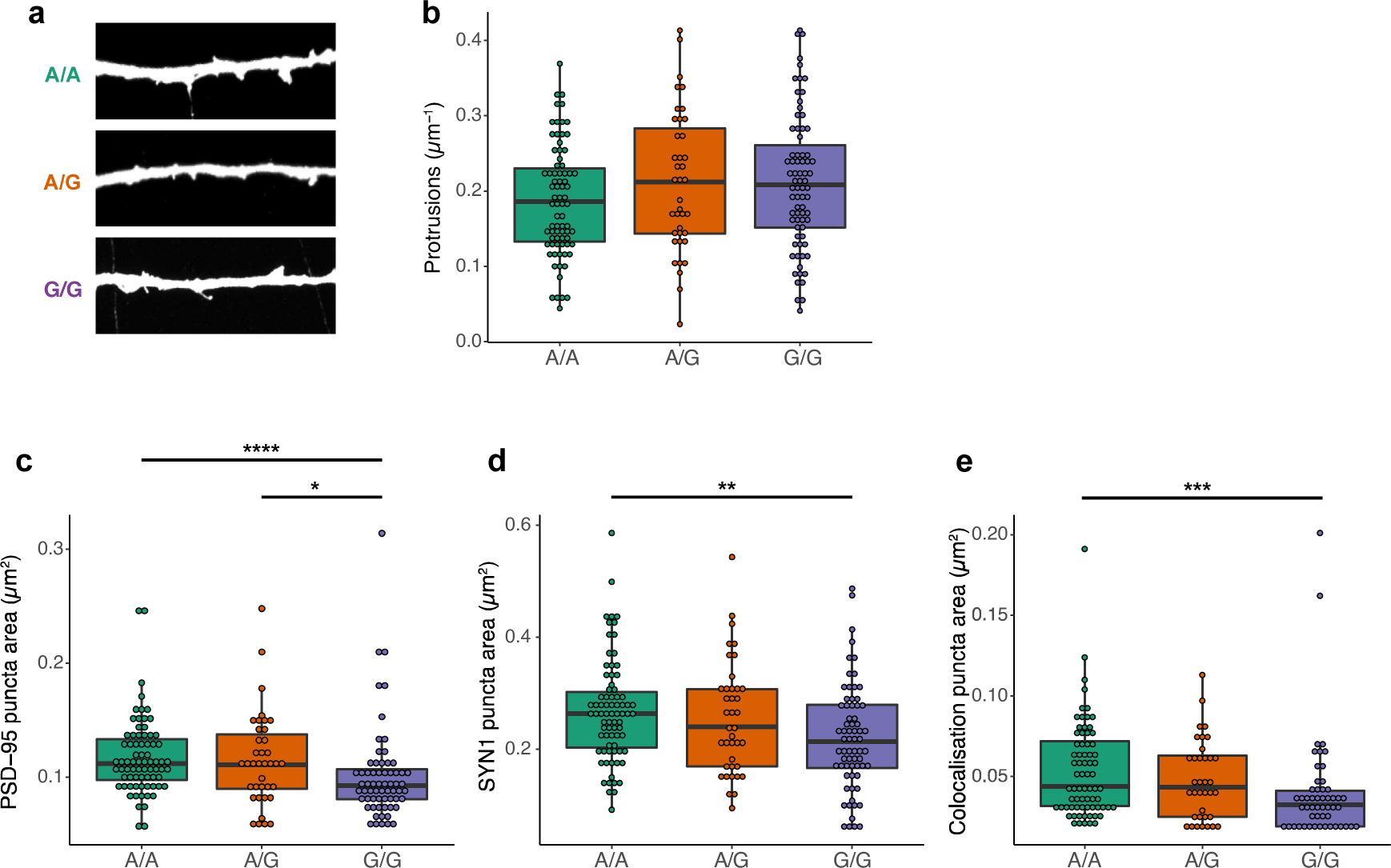
Protrusion density and puncta size analysis in NGN2-Glut of the rs2027349-edited isogenic lines. (a) Representative immunofluorescence images of dendrite and their associated neural protrusions in excitatory neurons. (b) Statistics of protrusion density in three genotypes, showing that genotype does not affect protrusion density. (c-e) Puncta size analysis. (c) Puncta size in different genotypes as measured by PSD-95 signal. (d) Puncta size in different genotypes as measured by SYN1 signal. (e) Puncta density in different genotypes as measured by the colocalization of PSD95:SYN1 signal, showing the A/A risk allele is associated with larger neural puncta size. In all cases, Kruskal-Wallis (non-parametric) test were used.

**Extended Data Fig. 8:**
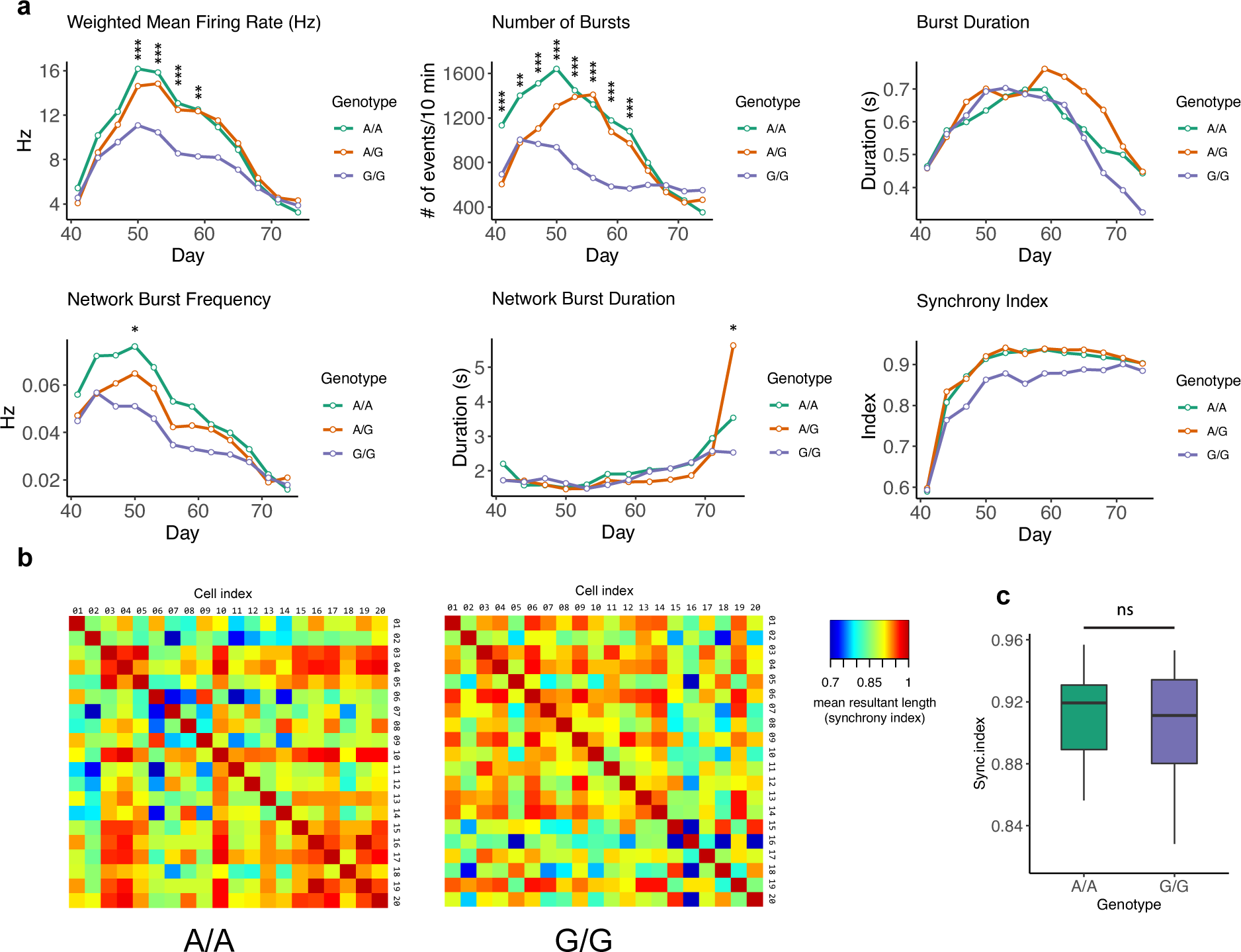
Additional data from MEA and Ca^2+^ imaging. (a) Full time-series (days 41-74) data from MEA assay of all six parameters measured. The significance level between A/A and G/G genotypes was indicated on the graphs. Two-way ANOVA with *post-hoc* correction for multiple testing was used. (b) Representative correlation matrix heatmap of neural activity synchrony index of rs2027349 A/A and G/G neurons. For each matrix heatmap, 20 neurons were randomly sampled from Ca^2+^ imaging data and the spike train distance model was applied on their detected peak series to generate the synchrony index heatmap (represented by the calculated mean resultant length). (c) Box-whisker plot of the correlation matrix in (b) with orthogonal elements removed, ns: not significant.

**Extended Data Fig. 9:**
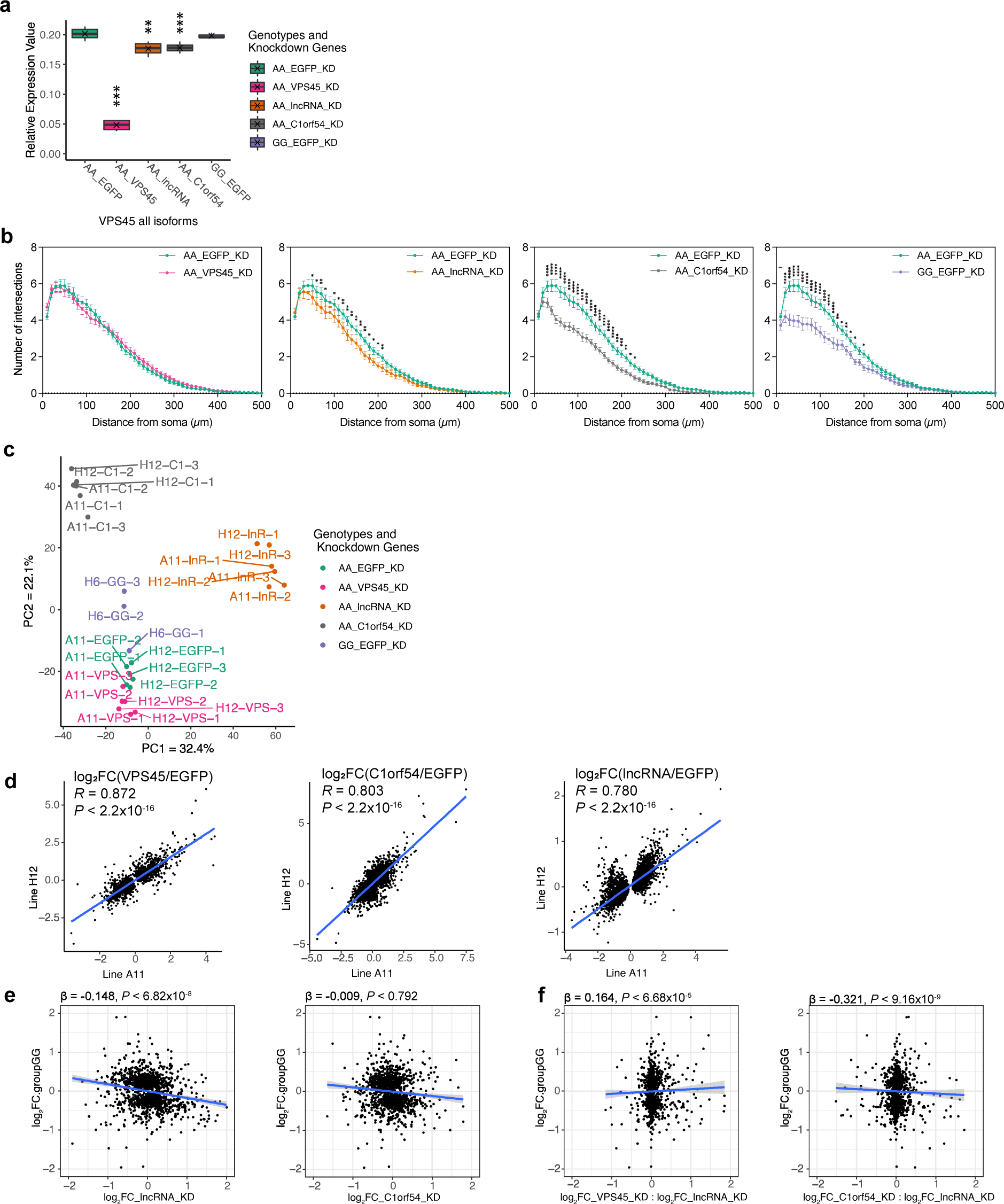
Additional data showing that *VPS45*, *AC244033.2*, and *C1orf54* interactively contribute to the altered neural phenotypes at the rs2027349 locus. (a) shRNA KD of any of the three targeted genes led to significant decrease in overall *VPS45* (all transcript isoforms) expression, with shRNA_*VPS45* showing most robust reduction of *VPS45*. Kruskal-Wallis (non-parametric) test with Dunn’s multiple comparisons and adjusted *P*-values. (b) Sample-wise split graph of Fig. 6, showing the Sholl analysis results of rs2027349 A/A line under different KD conditions in comparison with rs2027349 G/G line. Two-way ANOVA with *post-hoc* correction for multiple testing was used to test the significance of the differences. (c) Principal component analysis (PCA) plot of all RNA-seq samples in KD experiment showing the clustering of different rs2027349 A/A knockdown line and rs2027349 G/G control lines. (d) Scatter plot showing the strong correlation of shRNA KD effects between the two cell lines (A11 and H12). A union of genes found in at least one cell line were used. (e-f) Introducing interaction terms between the expression effects presents a marked improvement of model performance. Scatter plot showing the correlation and linear regression result between the log_2_ fold changes (FC) of DE genes in SNP-edited G/G neurons (G/G vs A/A) and the log_2_FC in different shRNA KD conditions in A/A neurons. (e) single-factor correlations, *AC244033.2* KD vs SNP editing; *C1orf54* KD vs SNP editing; (b) multi-factor correlations, *VPS45* KD x *AC244033.2* KD vs SNP editing; *C1orf54* KD x *AC244033.2* KD vs SNP editing.

**Extended Data Fig. 10:**
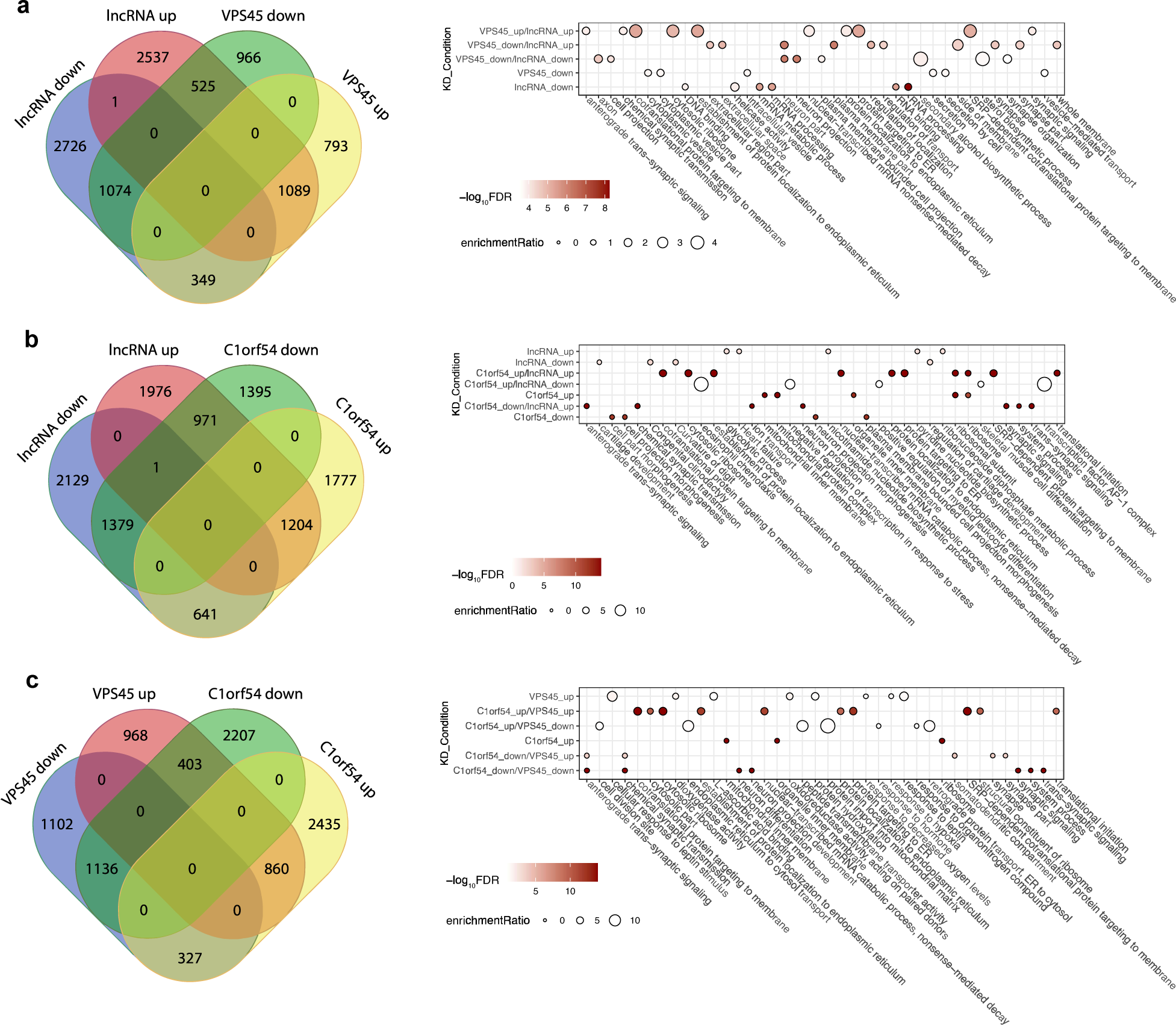
(a-c) Venn diagrams and bubble plots showing the overlap of gene counts between different KD conditions. In each figure, the right-panel bubble plot matches the left-panel Venn diagram. Only the top 40 GO terms (ranked by enrichment ratio) were shown in each bubble plot and only conditions (rows) with corresponding values in these terms were plotted. Circle size: enrichment ratio; fill: FDR.

**Extended Data Fig. 11:**
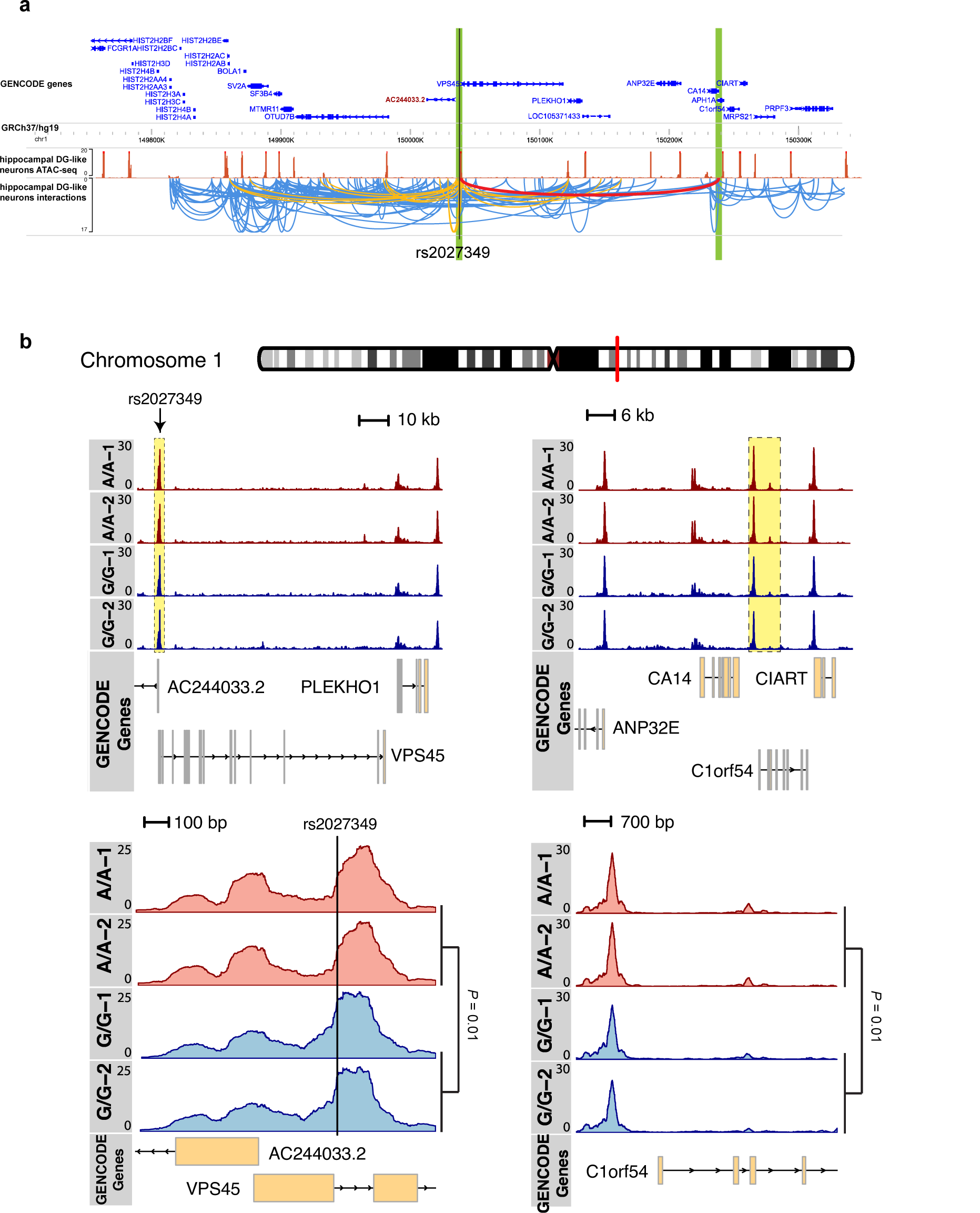
Long-range chromatin interactions and changes in local chromatin accessibility contribute to the cis-effect of rs2027349 on the expressions of multiple genes. (a) ATAC-seq and Capture Hi-C data from human brain tissue ^47^ showed that the rs2027349 locus has multiple long-range interactions (orange arcs) and, more specifically, connected to the promoter region of *C1orf54* at a distance (red arc and shaded regions). (b) CRISPR/Cas9-editing of rs2027349 led to subsequent alteration on both local and remote chromatin accessibility. Note the changes in normalized pile-up intensity on both the rs2027349 locus and regions proximal to *C1orf54* gene. Reads have been scaled and normalized in the unit of reads per genomic content (RPGC). Note that the lower read counts in A/A neurons at the rs2027349 site, which is consistent with the lower counts of A-containing ATAC-seq reads in A/G neurons in Fig. 1d. The lower counts of A-containing ATAC-seq reads at the SNP site were likely due to the stronger TF-binding protecting the TF-binding site from Tn5 accessibility ^13^.

## References

1. Consortium, T.S.W.G.o.t.P.G., Ripke, S., Walters, J.T. & O’Donovan, M.C. Mapping genomic loci prioritises genes and implicates synaptic biology in schizophrenia. medRxiv (2020).

2. Mullins, N. et al. Genome-wide association study of more than 40,000 bipolar disorder cases provides new insights into the underlying biology. Nat Genet 53, 817–829 (2021).

3. Demontis, D. et al. Discovery of the first genome-wide significant risk loci for attention deficit/hyperactivity disorder. Nat Genet 51, 63–75 (2019).

4. Grove, J. et al. Identification of common genetic risk variants for autism spectrum disorder. Nat Genet 51, 431–444 (2019).

5. Howard, D.M. et al. Genome-wide meta-analysis of depression identifies 102 independent variants and highlights the importance of the prefrontal brain regions. Nat Neurosci 22, 343–352 (2019).

6. Gandal, M.J. et al. Transcriptome-wide isoform-level dysregulation in ASD, schizophrenia, and bipolar disorder. Science 362(2018).

7. Rajarajan, P. et al. Neuron-specific signatures in the chromosomal connectome associated with schizophrenia risk. Science 362(2018).

8. Amiri, A. et al. Transcriptome and epigenome landscape of human cortical development modeled in organoids. Science 362(2018).

9. Li, M. et al. Integrative functional genomic analysis of human brain development and neuropsychiatric risks. Science 362(2018).

10. Nestler, E.J. & Hyman, S.E. Animal models of neuropsychiatric disorders. Nature Neuroscience 13, 1161–1169 (2010).

11. Wen, Z. et al. Synaptic dysregulation in a human iPS cell model of mental disorders. Nature 515, 414–8 (2014).

12. Schrode, N. et al. Synergistic effects of common schizophrenia risk variants. Nat Genet 51, 1475–1485 (2019).

13. Zhang, S. et al. Allele-specific open chromatin in human iPSC neurons elucidates functional disease variants. Science 369, 561–565 (2020).

14. Forrest, M.P. et al. Open Chromatin Profiling in hiPSC-Derived Neurons Prioritizes Functional Noncoding Psychiatric Risk Variants and Highlights Neurodevelopmental Loci. Cell Stem Cell 21, 305–318 e8 (2017).

15. Wang, D. et al. Comprehensive functional genomic resource and integrative model for the human brain. Science 362(2018).

16. de la Torre-Ubieta, L. et al. The Dynamic Landscape of Open Chromatin during Human Cortical Neurogenesis. Cell 172, 289–304 e18 (2018).

17. Consortium, G.T. et al. Genetic effects on gene expression across human tissues. Nature 550, 204–213 (2017).

18. Gutierrez-Arcelus, M. et al. Allele-specific expression changes dynamically during T cell activation in HLA and other autoimmune loci. Nat Genet 52, 247–253 (2020).

19. Buenrostro, J.D., Giresi, P.G., Zaba, L.C., Chang, H.Y. & Greenleaf, W.J. Transposition of native chromatin for fast and sensitive epigenomic profiling of open chromatin, DNA-binding proteins and nucleosome position. Nat Methods 10, 1213–8 (2013).

20. Zhang, Y. et al. Rapid single-step induction of functional neurons from human pluripotent stem cells. Neuron 78, 785–98 (2013).

21. Christian, K.M., Song, H. & Ming, G.L. Using Two- and Three-Dimensional Human iPSC Culture Systems to Model Psychiatric Disorders. Adv Neurobiol 25, 237–257 (2020).

22. De Los Angeles, A. et al. Induced Pluripotent Stem Cells in Psychiatry: An Overview and Critical Perspective. Biol Psychiatry 90, 362–372 (2021).

23. Townsley, K.G., Brennand, K.J. & Huckins, L.M. Massively parallel techniques for cataloguing the regulome of the human brain. Nat Neurosci 23, 1509–1521 (2020).

24. Fernando, M.B., Ahfeldt, T. & Brennand, K.J. Modeling the complex genetic architectures of brain disease. Nat Genet 52, 363–369 (2020).

25. Zhang, Y. et al. Model-based analysis of ChIP-Seq (MACS). Genome Biol 9, R137 (2008).

26. International Schizophrenia, C. et al. Common polygenic variation contributes to risk of schizophrenia and bipolar disorder. Nature 460, 748–52 (2009).

27. Schizophrenia Working Group of the Psychiatric Genomics, C. Biological insights from 108 schizophrenia-associated genetic loci. Nature 511, 421–7 (2014).

28. Datlinger, P. et al. Pooled CRISPR screening with single-cell transcriptome readout. Nat Methods 14, 297–301 (2017).

29. Xie, S., Duan, J., Li, B., Zhou, P. & Hon, G.C. Multiplexed Engineering and Analysis of Combinatorial Enhancer Activity in Single Cells. Mol Cell 66, 285–299 e5 (2017).

30. Yeo, N.C. et al. An enhanced CRISPR repressor for targeted mammalian gene regulation. Nat Methods 15, 611–616 (2018).

31. Dulubova, I. et al. How Tlg2p/syntaxin 16 ’snares’ Vps45. EMBO J 21, 3620–31 (2002).

32. McGuire, A.L. et al. The road ahead in genetics and genomics. Nat Rev Genet 21, 581–596 (2020).

33. Ang, C.E. et al. The novel lncRNA lnc-NR2F1 is pro-neurogenic and mutated in human neurodevelopmental disorders. Elife 8(2019).

34. Wang, J., Vasaikar, S., Shi, Z., Greer, M. & Zhang, B. WebGestalt 2017: a more comprehensive, powerful, flexible and interactive gene set enrichment analysis toolkit. Nucleic Acids Res 45, W130–W137 (2017).

35. Koopmans, F. et al. SynGO: An Evidence-Based, Expert-Curated Knowledge Base for the Synapse. Neuron 103, 217–234.e4 (2019).

36. Sharma, V. et al. eIF2alpha controls memory consolidation via excitatory and somatostatin neurons. Nature 586, 412–416 (2020).

37. Kirov, G. et al. De novo CNV analysis implicates specific abnormalities of postsynaptic signalling complexes in the pathogenesis of schizophrenia. Mol Psychiatry 17, 142–53 (2012).

38. Sey, N.Y.A. et al. A computational tool (H-MAGMA) for improved prediction of brain-disorder risk genes by incorporating brain chromatin interaction profiles. Nature Neuroscience 23, 583–593 (2020).

39. de Leeuw, C.A., Mooij, J.M., Heskes, T. & Posthuma, D. MAGMA: generalized gene-set analysis of GWAS data. PLoS Comput Biol 11, e1004219 (2015).

40. Nagel, M. et al. Meta-analysis of genome-wide association studies for neuroticism in 449,484 individuals identifies novel genetic loci and pathways. Nat Genet 50, 920–927 (2018).

41. Stahl, E.A. et al. Genome-wide association study identifies 30 loci associated with bipolar disorder. Nat Genet 51, 793–803 (2019).

42. Zhou, Z., Yuan, Q., Mash, D.C. & Goldman, D. Substance-specific and shared transcription and epigenetic changes in the human hippocampus chronically exposed to cocaine and alcohol. Proc Natl Acad Sci U S A 108, 6626–31 (2011).

43. Liu, J.Z. et al. Association analyses identify 38 susceptibility loci for inflammatory bowel disease and highlight shared genetic risk across populations. Nat Genet 47, 979–986 (2015).

44. Blizinsky, K.D. et al. Reversal of dendritic phenotypes in 16p11.2 microduplication mouse model neurons by pharmacological targeting of a network hub. Proc Natl Acad Sci U S A 113, 8520–5 (2016).

45. Yi, F. et al. Autism-associated SHANK3 haploinsufficiency causes Ih channelopathy in human neurons. Science 352, aaf2669 (2016).

46. Song, M. et al. Mapping cis-regulatory chromatin contacts in neural cells links neuropsychiatric disorder risk variants to target genes. Nat Genet 51, 1252–1262 (2019).

47. Wang, Y. et al. The 3D Genome Browser: a web-based browser for visualizing 3D genome organization and long-range chromatin interactions. Genome Biol 19, 151 (2018).

48. Penzes, P., Cahill, M.E., Jones, K.A., VanLeeuwen, J.E. & Woolfrey, K.M. Dendritic spine pathology in neuropsychiatric disorders. Nat Neurosci 14, 285–93 (2011).

49. Duan, J., Sanders, A.R. & Gejman, P.V. From Schizophrenia Genetics to Disease Biology: Harnessing New Concepts and Technologies. J Psychiatr Brain Sci 4(2019).

50. Ramocki, M.B. & Zoghbi, H.Y. Failure of neuronal homeostasis results in common neuropsychiatric phenotypes. Nature 455, 912–8 (2008).

51. Landek-Salgado, M.A., Faust, T.E. & Sawa, A. Molecular substrates of schizophrenia: homeostatic signaling to connectivity. Molecular Psychiatry 21, 10–28 (2016).

52. Javierre, B.M. et al. Lineage-Specific Genome Architecture Links Enhancers and Non-coding Disease Variants to Target Gene Promoters. Cell 167, 1369–1384 e19 (2016).

53. Li, X. & Fu, X.D. Chromatin-associated RNAs as facilitators of functional genomic interactions. Nat Rev Genet 20, 503–519 (2019).

54. Atianand, M.K. et al. A Long Noncoding RNA lincRNA-EPS Acts as a Transcriptional Brake to Restrain Inflammation. Cell 165, 1672–1685 (2016).

55. Allou, L. et al. Non-coding deletions identify Maenli lncRNA as a limb-specific En1 regulator. Nature 592, 93–98 (2021).

56. Pak, C. et al. Cross-platform validation of neurotransmitter release impairments in schizophrenia patient-derived NRXN1-mutant neurons. Proc Natl Acad Sci U S A 118(2021).

57. Dobin, A. et al. STAR: ultrafast universal RNA-seq aligner. Bioinformatics 29, 15–21 (2013).

58. Bray, N.L., Pimentel, H., Melsted, P. & Pachter, L. Near-optimal probabilistic RNA-seq quantification. Nat Biotechnol 34, 525–7 (2016).

59. Robinson, M.D., McCarthy, D.J. & Smyth, G.K. edgeR: a Bioconductor package for differential expression analysis of digital gene expression data. Bioinformatics 26, 139–40 (2010).

60. Xu, J. et al. Landscape of monoallelic DNA accessibility in mouse embryonic stem cells and neural progenitor cells. Nat Genet 49, 377–386 (2017).

61. van de Geijn, B., McVicker, G., Gilad, Y. & Pritchard, J.K. WASP: allele-specific software for robust molecular quantitative trait locus discovery. Nat Methods 12, 1061–3 (2015).

62. McKenna, A. et al. The Genome Analysis Toolkit: a MapReduce framework for analyzing next-generation DNA sequencing data. Genome Res 20, 1297–303 (2010).

63. Ramirez, F. et al. deepTools2: a next generation web server for deep-sequencing data analysis. Nucleic Acids Res 44, W160–5 (2016).

64. Bernstein, B.E. et al. The NIH Roadmap Epigenomics Mapping Consortium. Nat Biotechnol 28, 1045–8 (2010).

65. Liland, K.H., Almoy, T. & Mevik, B.H. Optimal choice of baseline correction for multivariate calibration of spectra. Appl Spectrosc 64, 1007–16 (2010).

66. Liao, Y., Smyth, G.K. & Shi, W. The R package Rsubread is easier, faster, cheaper and better for alignment and quantification of RNA sequencing reads. Nucleic Acids Res (2019).

67. Harrow, J. et al. GENCODE: the reference human genome annotation for The ENCODE Project. Genome Res 22, 1760–74 (2012).

68. Mayfield, R.D., Harris, R.A. & Schuckit, M.A. Genetic factors influencing alcohol dependence. Br J Pharmacol 154, 275–87 (2008).

69. Gaiteri, C. & Sibille, E. Differentially expressed genes in major depression reside on the periphery of resilient gene coexpression networks. Front Neurosci 5, 95 (2011).

70. Sey, N.Y.A. et al. A computational tool (H-MAGMA) for improved prediction of brain-disorder risk genes by incorporating brain chromatin interaction profiles. Nat Neurosci 23, 583–593 (2020).

71. Koopmans, F. et al. SynGO: An Evidence-Based, Expert-Curated Knowledge Base for the Synapse. Neuron 103, 217–234 e4 (2019).

72. Pardinas, A.F. et al. Common schizophrenia alleles are enriched in mutation-intolerant genes and in regions under strong background selection. Nat Genet 50, 381–389 (2018).

73. Bayes, A. et al. Characterization of the proteome, diseases and evolution of the human postsynaptic density. Nat Neurosci 14, 19–21 (2011).

74. Victor, J.D. & Purpura, K.P. Nature and precision of temporal coding in visual cortex: a metric-space analysis. J Neurophysiol 76, 1310–26 (1996).

